# An interpretable deep learning framework for classifying neuronal morphologies using topology and graph neural networks

**DOI:** 10.1101/2024.09.13.612635

**Authors:** Lida Kanari, Stanislav Schmidt, Francesco Casalegno, Emilie Delattre, Jelena Banjac Lukic, Thomas Negrello, Ying Shi, Julie Meystre, Michaël Defferrard, Felix Schürmann, Henry Markram

## Abstract

Neuronal shape determines how neurons process and integrate information, yet a consistent and objective classification of neuronal morphologies remains elusive. Current approaches rely heavily on subjective expert views or on predefined features, limiting reproducibility and interpretability. Here, we present an interpretable deep learning framework that unifies topological data analysis, graph neural networks, and traditional morphometrics to classify neuronal morphologies objectively and transparently. Our framework compares complementary mathematical representations of neurons to capture geometric, topological, and graph-structural information. Then it benchmarks their performance against expert-labeled datasets. We show that topology- and graph-based models achieve accuracies comparable to human experts, revealing that both global branching invariants and local connectivity patterns are essential to define morphological cell types. Using explainable artificial intelligence methods, we identify structural features driving each classification decision, bridging computational and neuroanatomical interpretations. This open source and reproducible approach provides a foundation for scalable, interpretable and biologically meaningful neuronal taxonomy, enabling consistent comparisons between data sets and species.

## 1 Introduction

Neurons can be broadly categorized into morphologically and functionally distinct classes, such as excitatory pyramidal neurons and inhibitory interneurons, which together form cortical circuits supporting brain computations (Häusser et al., 2000; Schaefer et al., 2003; Chklovskii, 2004). Pyramidal neurons, which constitute the majority of cortical neurons (Spruston, 2008; Markram et al., 2015), are characterized by their triangular soma and extensive dendritic arbors and exhibit highly diverse dendritic and axonal shapes that determine their roles in cognition and perception (Häusser et al., 2000). Pyramidal cells define the excitatory output of the cortex and are central to information integration and long-range signaling (Häusser et al., 2000), while interneurons support inhibitory modulation of local cortical circuitry (Häusser et al., 2000; Schaefer et al., 2003; Yamahachi et al., 2009). Importantly, differences in neuronal morphology are closely related to functional specialization, since structural features such as dendritic branching patterns (Kanari et al., 2019) and axonal projections (Häusser et al., 2000; Schaefer et al., 2003) determine how neurons integrate inputs and contribute to circuit-level computations and behavior (Marcassa et al., 2024). A major goal for neuroscience is to establish a robust and universal framework for classifying neurons into morphological classes (Ascoli et al., 2008; DeFelipe et al., 2013; Kanari et al., 2019). This task has gained new momentum with the growing availability of standardized reconstructions from large-scale public resources such as neuromorpho.org (Ascoli et al., 2007) and the Allen Brain Atlas (Gouwens et al., 2019).

Defining a state-of-the-art neuronal classification remains challenging, as neurons can be grouped according to different structural, functional, or multimodal perspectives. Traditional techniques based on morphometrics (Ascoli et al., 2008; DeFelipe et al., 2013; Gouwens et al., 2019) have been complemented by topological approaches (Li et al., 2017; Kanari et al., 2018, 2019; Laturnus et al., 2020; Khalil et al., 2022) and by recent advances in machine learning, including graph neural networks (GNNs) (Henaff et al., 2015; Defferrard et al., 2017; Weis et al., 2025). More recently, multimodal classification frameworks have sought to integrate morphological, electrical, and transcriptomic information (Gouwens et al., 2019; Scala et al., 2020). In the present work, we focus exclusively on morphology to examine the relative performance and interpretability of different classification schemes under the realistic constraint of limited expert reproducibility (DeFelipe et al., 2013).

A key question motivating this study is *what can we learn from a classification scheme beyond accuracy itself?* To answer this, we systematically compare a range of morphological representations and learning strategies to understand how distinct cell types differ in shape and structure. Crucially, we argue that any attempt to improve classification performance must be interpreted in light of the inherent uncertainty of expert-defined labels, as demonstrated by inter-rater variability (DeFelipe et al., 2013). Our goal is therefore not to propose a novel classifier that supersedes existing methods, but to develop a comprehensive and mathematically robust framework for understanding morphological diversity through multiple, interpretable representations.

Traditionally, morphological classification has relied on single-valued morphometrics such as total length, tortuosity, or tree asymmetry (Ascoli et al., 2008). Although informative, these one-dimensional projections lose substantial information about the complex geometry of neuronal arbors. To capture this complexity, we combine several complementary approaches: (i) traditional morphometrics (Ascoli et al., 2008), (ii) the Topological Morphology Descriptor (TMD) (Kanari et al., 2018), which encodes the branching structure into a topological descriptor, and (iii) graph neural networks (GNNs) (Defferrard et al., 2017; Weis et al., 2025), which operate directly on neuronal graphs. These representations bridge complementary levels of description: morphometrics capture measurable geometric quantities, TMD quantifies global topological invariants, and GNNs learn hierarchical graph features from the raw structure itself.

Using a diverse dataset of cortical neuronal morphologies (Markram et al., 2015; Muralidhar et al., 2013; Winnubst et al., 2019), comprising pyramidal neurons and interneurons, we perform classification using graph neural networks (GNNs), topological morphology descriptors (TMD), and classical morphometric features, and systematically evaluate the performance of these methodologies across heterogeneous data. We show that advances in machine learning eliminate the need for manual feature selection (Markram et al., 2015; Laturnus et al., 2020; Scala et al., 2020). By combining XGBoost with a comprehensive library of morphometrics, a superset of previously optimized feature sets (DeFelipe et al., 2013; Markram et al., 2015; Laturnus et al., 2020; Scala et al., 2020), we achieve higher accuracy than with any hand-picked subsets. We then compare these results to those obtained using TMD-based classifiers and GNNs. Neuronal morphologies are represented as directed adjacency matrices for GNNs, similar to the unsupervised learning framework proposed in (Weis et al., 2021).

Recent classification studies emphasize maximizing accuracy against expert-defined labels (Laturnus et al., 2020; Weis et al., 2021), but these labels themselves are not entirely consistent (DeFelipe et al., 2013). Human experts apply variable and evolving criteria, leading to limited inter- and intra-rater reliability. Accounting for this variability is essential for evaluating how meaningful is a specific classifier. For example, a classifier that performs with high accuracy on random labels is not improving our understanding of neuronal morphologies. Our analysis includes datasets independently labeled by two experts, allowing us to quantify inter-rater agreement and to use this as a principled bound for meaningful classification performance. In addition, our results are compared with a baseline of classification on a randomized set of labels to indicate baseline accuracy.

Our results demonstrate that no single classifier is universally optimal. Instead, different methods capture distinct aspects of neuronal structure: morphometrics are optimal for interneurons, while topological and graph-based approaches perform best for pyramidal cells. Importantly, the best-performing models reach accuracy levels comparable to inter-rater agreement, establishing this agreement as a realistic benchmark for neuronal morphology classification. We thus propose a unified and interpretable framework that integrates morphometric, topological, and graph-based analyses for robust and reproducible classification of neuronal morphologies. The proposed metholodogy can be applied to any dataset for which neurons are reconstructed with sufficient detail, such as the Microns (The MICrONS Consortium et al., 2025), Harvard (Shapson-Coe et al., 2024) or FlyWire (Dorkenwald et al., 2020) datasets.

All code and examples are available in our open-source repository at https://github.com/BlueBrain/morphoclass, enabling the application of our framework to new datasets and classification tasks.

## 2 Results

We combined different feature extractions for an increasing level of complexity (one-dimensional morphometrics (Ascoli et al., 2007), deepwalk (Perozzi et al., 2014), see DeepWalk, TMD (Kanari et al., 2018), see TMD, and morphology graph) with a variety of machine learning techniques (traditional machine learning models such as decision trees and XG-Boost (Pedregosa et al., 2011a), see Traditional Machine Learning Models, convolutional neural networks, see Convolutional Neural Networks, PersLay (Carriere et al., 2020), see A PersLay-Based Classifier and graph neural networks (Henaff et al., 2015; Defferrard et al., 2017), see Graph Neural Networks) and evaluated their classification accuracy. The combination of features and machine learning techniques will be referred to as “classification methods” throughout this paper (Figure 1).

**Figure 1:**
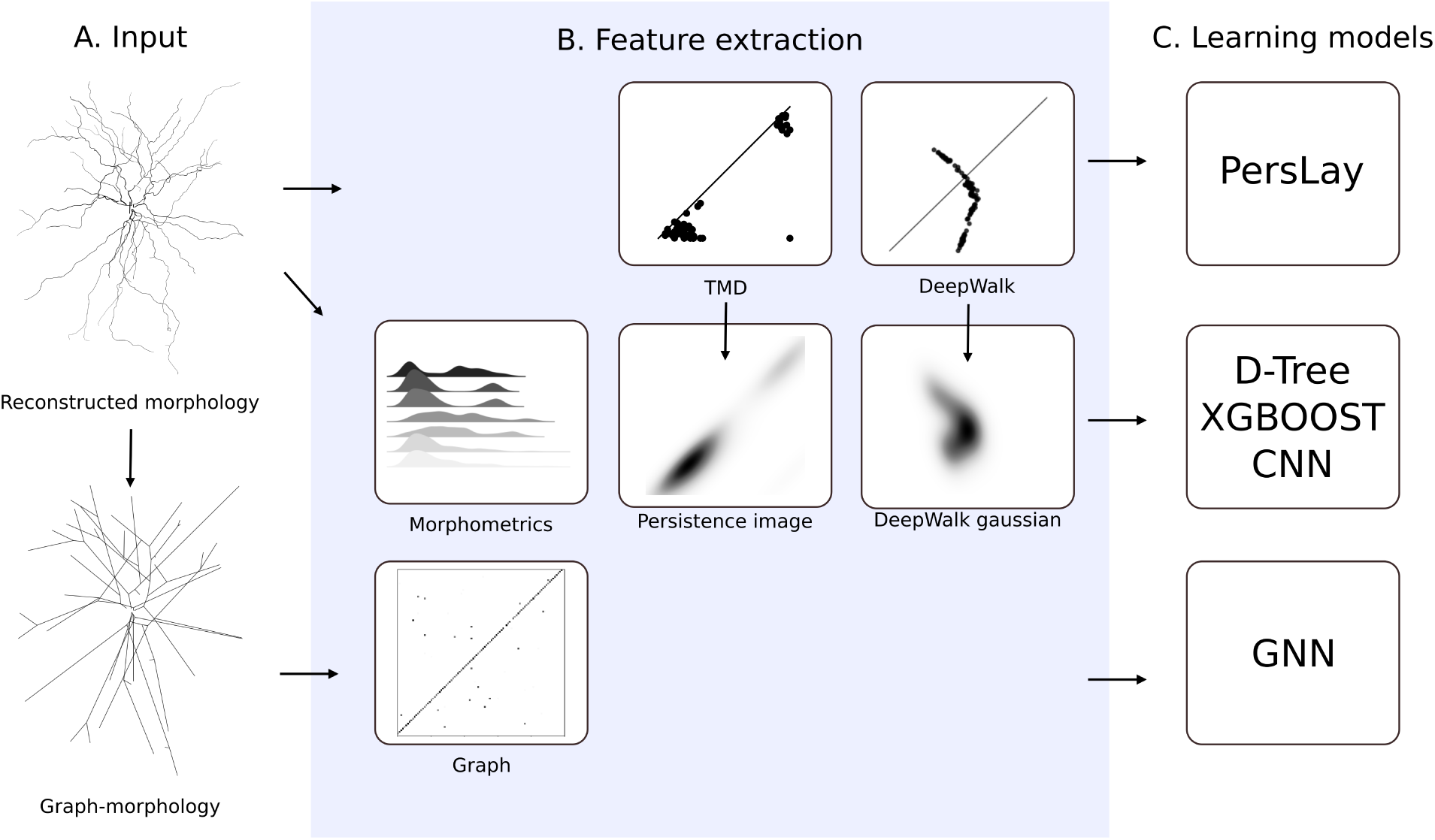
Classification method overview. The neuronal morphology (A), either as a reconstructed morphology or a simplified graph is used as input. The feature extraction step (B) reduces the dimensionality of the data either by vectorization (Morphometrics, TMD, DeepWalk) or by graph embedding to be used as inputs in a variety of machine learning techniques. (C) The learning models (PesrLAY, Decision tree, XGBoost, CNN, GNN) output the model accuracy.

The performance of each classification method was evaluated (see Evaluation metrics) on two primary datasets, each separated by the cortical layer: rodent interneurons and pyramidal cells. These datasets have been evaluated independently in the past with conflicting results (DeFelipe et al., 2013; Kanari et al., 2019; Laturnus et al., 2020). However, the morphologies of interneurons and pyramidal cells present different challenges. For example, interneurons are typically categorized based on their axonal projections, while pyramidal cells differ mainly on the morphology of apical dendrites. Therefore, evaluating the capabilities of different classifiers with multiple datasets is important. The datasets we used are described in detail below:

- **Interneurons LNMC**: 583 interneurons from layers 1 to 6 were collected in the LNMC laboratory (Markram et al., 2015; Muralidhar et al., 2013). Patch-clamped neurons were filled with biocytin and reconstructed by experts. The expert classification was performed to assign expert labels to each cell.
- **Pyramidal cells LNMC**: 465 pyramidal cells from layers 2 to 6 were collected in the LNMC laboratory (Markram et al., 2015). Patch-clamped neurons were filled with biocytin and reconstructed by experts. The expert classification was performed to assign expert labels to each cell.
- **Pyramidal cells Janelia**: 58 pyramidal cells of layer 5 were stained in fluorescent dye and reconstructed semi-automatically in the Janelia laboratory (Winnubst et al., 2019). The expert classification was performed to assign expert labels to each cell.

First, the accuracy of different classifiers based on a variety of features was computed for each of the datasets above. The performance of different methods was evaluated based on accuracy (see Methods: Evaluation metrics). Alternative accuracy methods (balanced accuracy, F1 score: micro, macro, and weighted) were computed, but the results were quantitatively, and not qualitatively different. Therefore, we chose to present the accuracy for compatibility with the previous literature (Kanari et al., 2019; Laturnus et al., 2020).

### 2.1 Comparison of classification methods

We investigated whether a single classification method could consistently outperform all others across the chosen datasets. We found that this is not the case for the neuronal morphology datasets. Due to the complexity of the experimental data, the small sample size, and the noise in the data, it was not possible to identify a single classification method that consistently outperforms the others. Consequently, we present a concise summary of our findings (Figure 2). Note that although we assessed numerous classification methods, including PersLay (Carriere et al., 2020), decision trees and random forests, and different combinations of features and classifiers, only a subset is presented in the main results of the paper. Specifically, we include the top-performing classifiers and at least one classifier for each feature extraction method (morphometrics, topology, graph, deepwalk). The summarized results are detailed in the following sections.

**Figure 2:**
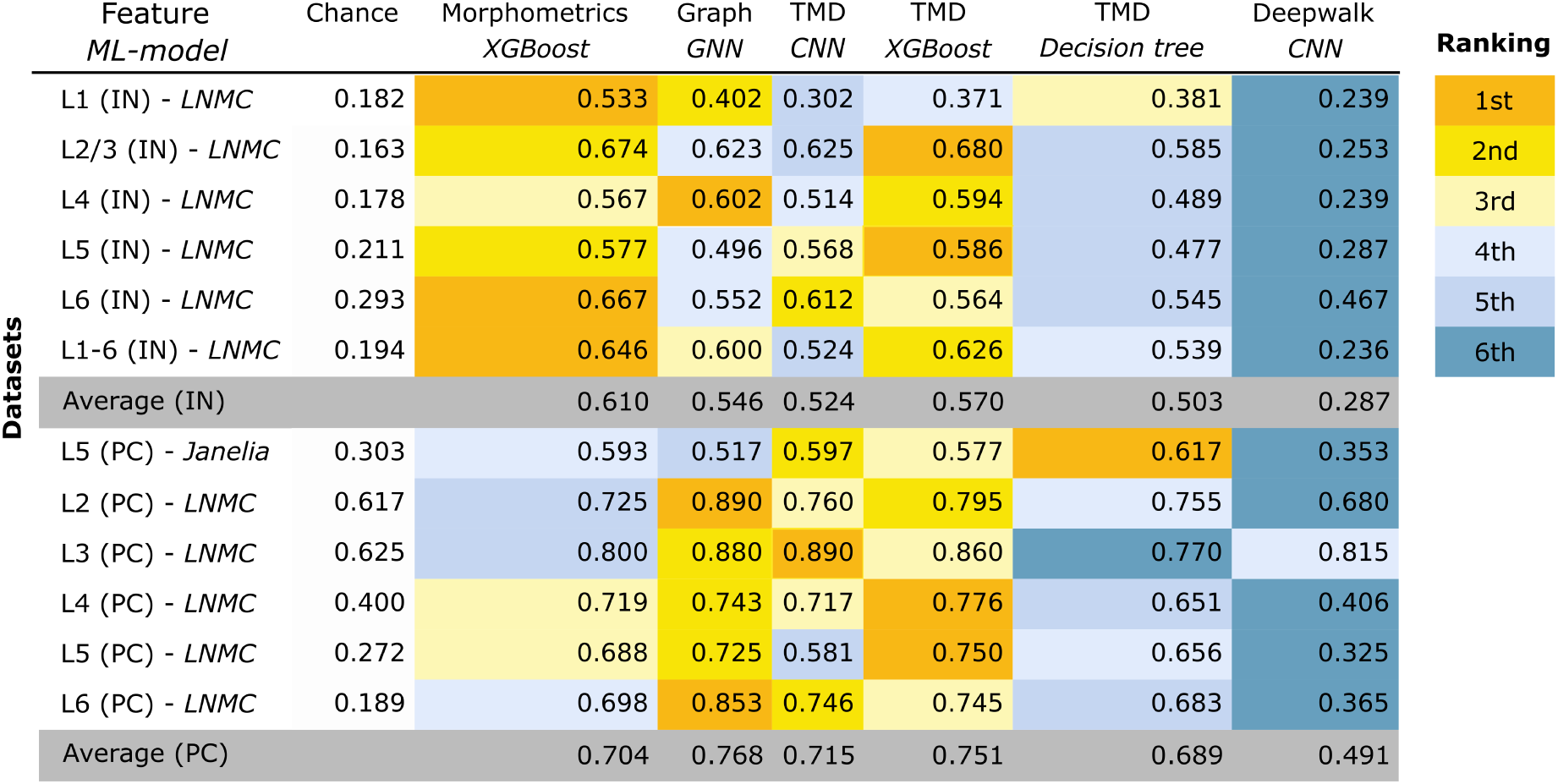
Table of classification accuracy (unbalanced) across different methods. Each column corresponds to a specific classification method, defined by a combination of feature representation (morphometrics, graph-based, or TMD-based) and learning algorithm (XGBoost, GNN, decision tree, CNN). Rows represent individual datasets for interneurons (IN) and pyramidal cells (PC), with average accuracies reported for each group. The first column indicates dataset-specific chance levels. Within each row, color coding reflects relative performance ranking across methods, from best (orange) to worst (blue).

The highest classification accuracy, ≈ 90%, was achieved by two techniques: *GNN* and *TMD-CNN*, specifically on the superficial pyramidal cell dataset (cortical layers 2 and 3 accordingly). However, this trend was not consistently observed in the interneuron dataset. For interneurons stratified per cortical layer (L1-L6), a lower performance was typically observed for all methods.

We observe that in all cases, the *deepwalk* feature consistently performed worse than all other methods. Therefore, deepwalk is not appropriate for discriminating neuronal morphologies. The discrepancy between accuracy scores per dataset for the same classification method, ranging from 50% to 90%, can be attributed to the high variability and the unbalanced nature of the biological datasets, which include classes between 5 and 50 cells (see details in sections Section 2.2.1, Section 2.3).

To further assess the performance of the classification methods, we provide confusion matrices (Figure S4) for the top three classifiers (*GNN*, *TMD-CNN*, and *morphometrics*) and the one with the lowest score *deepwalk*. The confusion matrix visually compares predicted labels to true labels for each class, with perfect accuracy represented by a diagonal where all predicted labels align with true labels. In the case of the three top-performing classifiers (GNN - graph, CNN - TMD and morphometrics), a distinct diagonal pattern is evident. On the contrary, for the lowest-performing classifier (deepwalk), the confusion matrices show a lack of clear diagonal separation, consistent with the lower accuracy scores. For the rest of the analysis, we present the results of the three top-performing classification methods for the different datasets.

#### 2.1.1 Optimal feature selection for morphometrics

We also experimented with alternative sets of morphometrics including subsets of morphometrics as presented in (Ascoli et al., 2007; Gouwens et al., 2019; Laturnus et al., 2020). The superset of 52 tree morphometrics (Table S1) per neurite type (axon, basal, and apical dendrites) and 12 for overall morphology (Table S2) outperformed any other subset. Therefore, we conclude that there is no optimal manual selection of morphometrics and it is more efficient to present the largest possible set of features as input to the XGBoost classifier for optimal results.

#### 2.1.2 Complementary of TMD - GNN based methods

We computed per-sample predictions for the different methods and summarized the results for all pyramidal cells and interneuron classes for GNN - graph, CNN - TMD and morphometrics Figure 3. Across datasets, TMD and GNN exhibit substantial overlap in correct predictions, while also capturing distinct subsets of samples, indicating complementary strengths rather than a single consistently superior method. TMD and GNN generally outperform morphometrics, which show fewer uniquely correct predictions and a higher proportion of misclassified samples.

**Figure 3:**
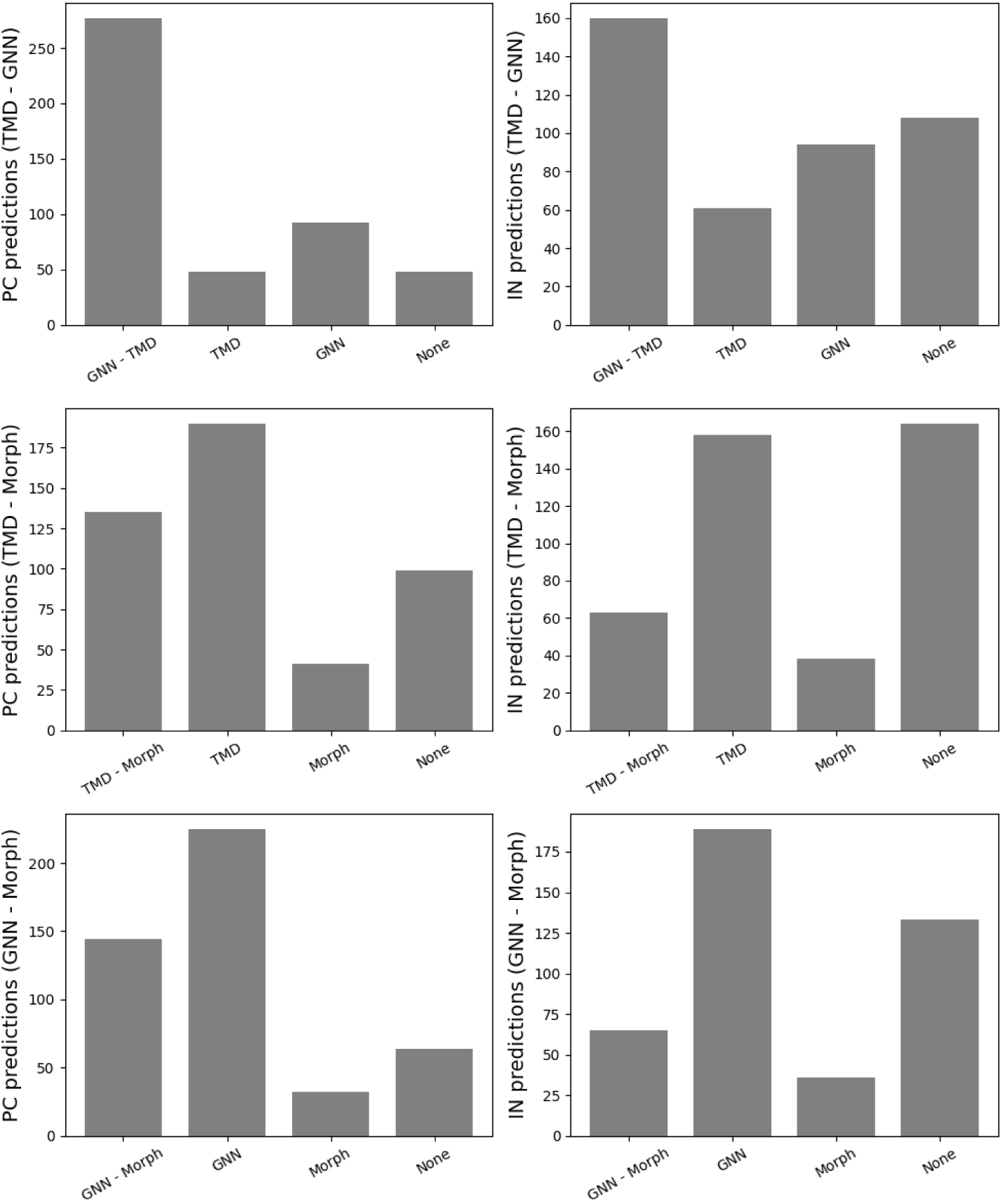
Comparison of classification performance across TMD, GNN, and morphometric approaches. Bar plots show the distribution of predictions for pyramidal cell (PC) and interneuron (IN) datasets, partitioned into samples correctly classified by both methods, only one method, or neither. For each pairwise comparison (TMD vs GNN, TMD vs Morphometrics, GNN vs Morphometrics), the number of correctly classified samples highlights differences in predictive behavior. Across datasets, TMD and GNN exhibit substantial overlap in correct predictions, while also capturing distinct subsets of samples, indicating complementary strengths rather than a single consistently superior method. TMD and GNN outperform morphometrics, which show fewer uniquely correct predictions and a higher proportion of misclassified samples.

To evaluate whether topology-based (TMD) and graph-based (GNN) approaches capture complementary information, we quantified the extent to which their predictions and representations overlap. Specifically, we analyzed per-sample prediction agreement in Layer 5 pyramidal cells and observed a substantial fraction of cases where only one of the two methods produced the correct classification, indicating non-overlapping error modes (Figure S 7).

In addition, we analyzed the non-overlaping errors for layer 5 pyramidal cells per class (see supplementary Figure S 8). The results reveal a class-dependent complementarity between the two approaches: GNN achieves higher unique performance for classes L5 TPC:B, L5 TPC:C, and L5 UPC. In contrast, TMD-CNN shows higher performance for L5 TPC:A, which is the most abundant class of layer 5 pyramidal cells.

These results indicate that different morphological features are preferentially captured by graph-based and topological representations, providing further evidence that the two methods encode distinct and complementary aspects of neuronal structure, rather than merely exhibiting dataset-dependent performance differences.

### 2.2 Classification of pyramidal cells

In this section, we focused on the dendritic reconstructions of 465 pyramidal cells from layers 2 to 6 (Figure 4A-B) (Markram et al., 2015; Kanari et al., 2019). The cell types of pyramidal cells are described in detail in Table S3. The highest classification accuracy was achieved by two techniques: graph neural networks (GNN) and convolutional neural networks (CNN) on TMD images. These methods outperformed others across various evaluation metrics. An analysis of variance (ANOVA) was performed to test the statistical significance of our results (see SI:ANOVA). The ANOVA results (see SI:ANOVA) revealed significant differences in performance between the methods, demonstrating higher performance in *GNN* and *TMD-CNN* classifiers, followed by *morphometrics*.

**Figure 4:**
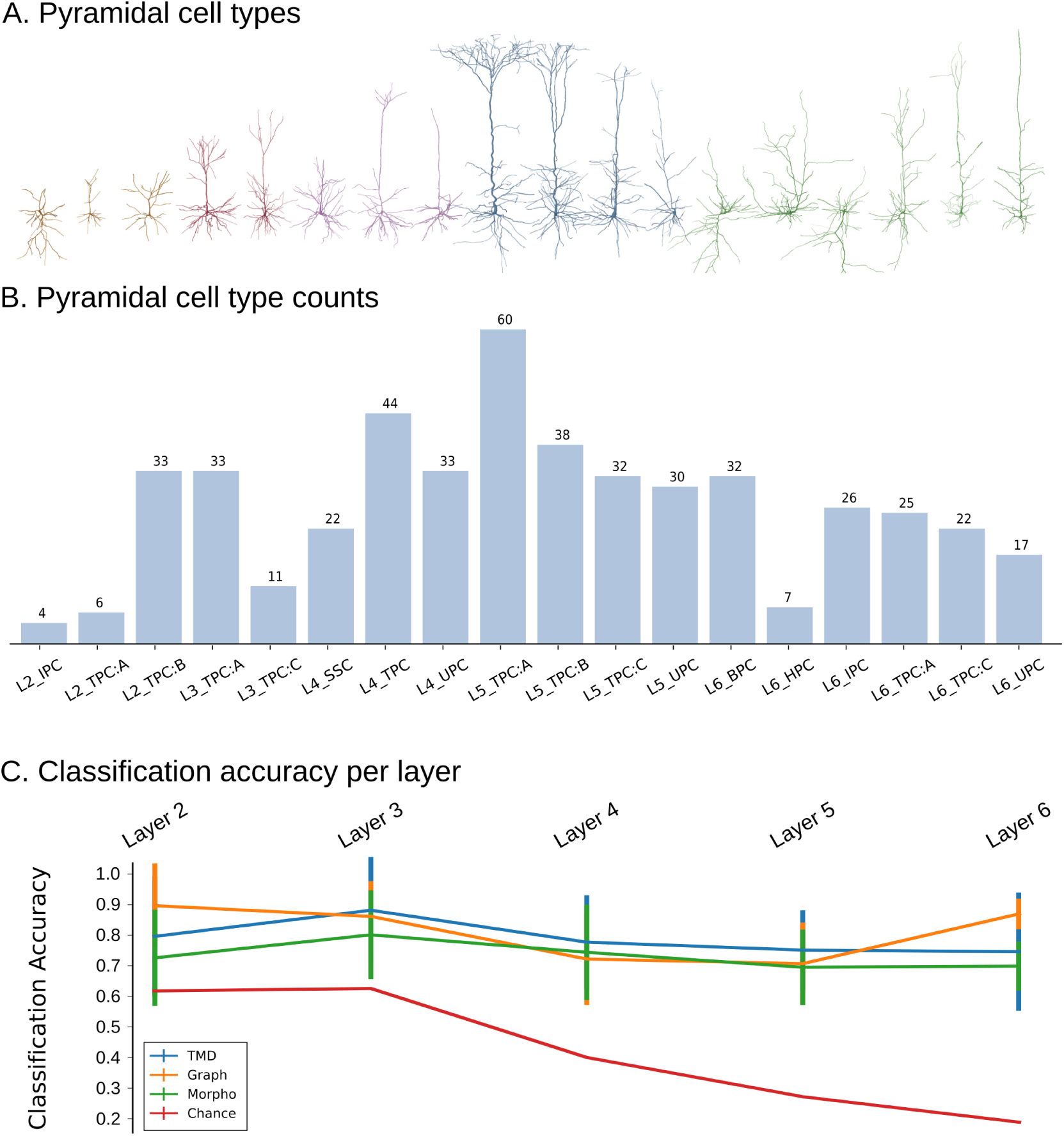
Pyramidal cell types, dataset composition, and classification performance. (A) Representative examples of expert-defined pyramidal cell types corresponding to the classes shown in (B). Colors indicate cortical layers: layer 2 (yellow), layer 3 (red), layer 4 (purple), layer 5 (blue), and layer 6 (green). (B) Number of cells per pyramidal cell type (see also Table S3). (C) Classification accuracy across cortical layers for different methods: TMD-based models (XGBoost), graph-based models (GNN), morphometric features, and chance level. Results are aggregated per layer, highlighting differences in performance across representations.

A comprehensive analysis revealed that features based on apical dendrites consistently outperformed features based on axons or basal dendrites (see Table S5). Overall, the classification of pyramidal cells yielded well-defined and stable classes, which could be reliably identified and supported by multiple classification methods. This suggests the presence of distinct morphological characteristics that distinguish different types of pyramidal cells and can be supported by GNN and TMD. The differences between these classes are further discussed in Explainable artificial intelligence: XAI.

#### 2.2.1 Comparison to inter-rater expert agreement

The classification of pyramidal cells from layer 5 (Janelia) was evaluated by two individual experts to compute the inter-rater agreement accuracy (see Inter-rater agreement), which was calculated at ≈ 80%. Most classification methods, including GNN, and TMD-CNN, were able to achieve accuracies comparable to the agreement between human raters. The mean accuracy for all BBP PCs datasets is (81%±7%) for GNN, (74%±9%) for TMD-CNN, (79% ± 4%) for TMD-XGBoost and (72% ± 4%) for morphometrics-XGBoost. This indicates that the GNN and TMD-XGBoost classification methods produce consistent and reliable results (around 80%), consistent with stable morphological classes. For Janelia dataset, the accuracy is smaller for all classification methods due to the small sample size. In addition, the classification accuracy was compared to the chance levels (Figure 2 and Figure 4C), as described in Section 4.6.2.

### 2.3 Classification of interneurons

The dataset focused on differentiation based on 583 interneurons from layers 1 to 6 (Figure 5A-B, Figure S5) (Markram et al., 2015; Muralidhar et al., 2013). The cell types of interneurons are described in detail in Table S4. A comprehensive analysis revealed that axonal-based features contributed more significantly in the classification accuracy compared to dendrites (see Table S5). Interestingly, unlike pyramidal cells, using a combination of basal dendrites and axons, improved the accuracy of interneuron classification, but not to statistically significant levels (see Table S5). In addition, we found that axonal features were more significant for the classification decision, compared to dendritic features (see Figure S2). Therefore, the classification results presented focus on axonal features for TMD-CNN and GNN.

**Figure 5:**
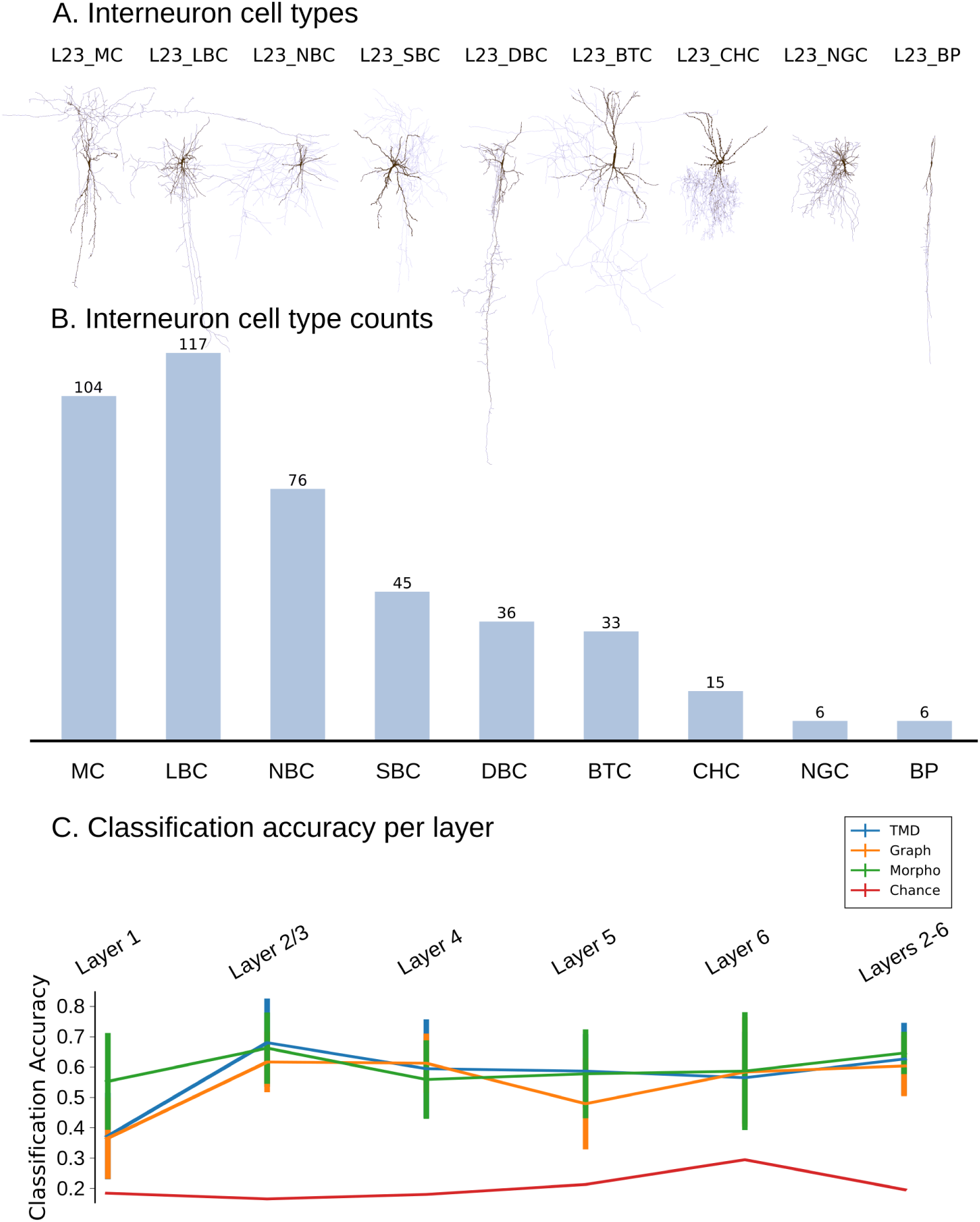
Interneuron cell types, dataset composition, and classification performance. (A) Representative examples of expert-defined interneuron cell types corresponding to the classes shown in (B). (B) Number of cells per pyramidal cell type (see also Table S4). (C) Classification accuracy across cortical layers for different methods: TMD-based models (XGBoost), graph-based models (GNN), morphometric features, and chance level. Results are aggregated per layer, high-lighting differences in performance across representations.

For the classification of interneurons stratified per cortical layers (1-6), all classifiers performed poorly. We hypothesize that this is due to the large number of classes and the small sample sizes. By merging interneurons of layers 2–6, the classification accuracy improved across all classification methods, reaching 65%. However, classification accuracy remained overall low compared to the classification accuracy of pyramidal cells, a result that generalizes across classifiers. Our findings suggest that interneurons in the cortex should be studied across layers rather than separated by layers. This is due to the smaller extend of interneurons, which constrains most cell types within the boundaries of a cortical layer. As a result, the difference in interneurons across layers is smaller than the difference across cell types. This holistic approach allows for a more accurate classification and understanding of interneuron diversity within the cortical microcircuit.

### 2.4 Explainable artificial intelligence: XAI

One of the main advantages of GNN and CNN is that they can be used to propose an explanation of the observed accuracy scores. Explainable artificial intelligence (XAI) is essential to interpret the results of classifiers and understand the differences between cell types in a more comprehensive manner. The use of Grad-CAM (Gradient-weighted Class Activation Mapping (Selvaraju et al., 2016)) as an XAI (Zhou et al., 2015; Lipton, 2016; Adadi and Berrada, 2018) provides valuable insights into the decision-making process of both CNN and GNN models (see Explainable Artificial Intelligence (XAI)). However, it is important to note that the interpretation of Grad-CAM differs between CNN and GNN methodologies due to their distinct input representations. For CNNs, which operate on image data, Grad-CAM highlights image regions of high importance for the classification decision, whereas for GNNs, which operate on graph-structured data, it identifies influential nodes in the graph structure (see Explainable Artificial Intelligence (XAI)).

#### 2.4.1 XAI: single-cell analysis

To gain insight into the decision-making processes of the models at the level of individual neurons, we applied Grad-CAM to both CNN–TMD and Graph–GNN classifiers (Figures S 9, S 10), using as an example the classification of layer 5 pyramidal cells. These approaches provide complementary perspectives by projecting model attention onto two distinct representations: the persistence image for the CNN–TMD and the neuronal graph for the GNN.

In the CNN–TMD framework (Figure S 9, Figure 6A), Grad-CAM identifies regions of the persistence image that contribute most strongly to the classification. Across correctly classified samples, these regions consistently coincide with areas of high density in the persistence representation, indicating that the model relies on robust topological features. Even at the level of individual cells, the highlighted regions closely resemble the dominant structures observed in class-average representations, suggesting that the model captures stable global signatures of neuronal morphology.

**Figure 6:**
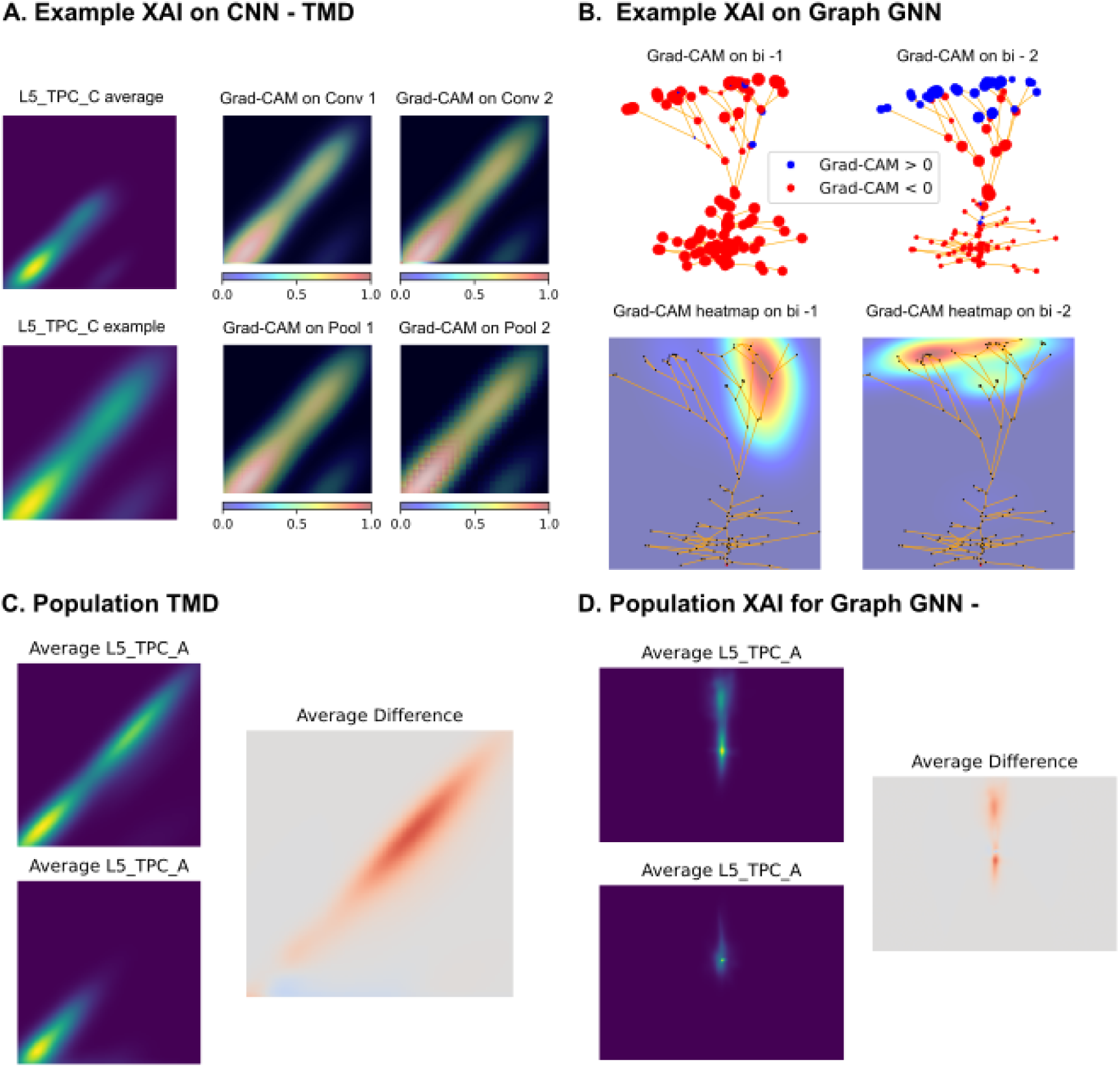
Grad-CAM-based explainability for CNN-TMD and Graph-GNN. (A) Example of a CNN–TMD model correctly classifying a Layer 5 TPC C pyramidal cell. On top we show the class-average TMD representation, on bottom an individual example. Grad-CAM highlights across convolutional and pooling layers (Conv1, Conv2, Pool1, Pool2) focus on the shape of the persistence images. (B) Example of a Graph–GNN model correctly classifying a Layer 5 pyramidal cell. Node-level Grad-CAM scores (top) and the corresponding spatial heatmaps (bottom) for the two layers (b1-1, bi-2). (C) Averaged TMD based persistence images across all the pyramidal cells in two classes L5 TPC A and L5 TPC C, and their respective average difference. (D) Averaged Grad-CAM heatmaps across all the pyramidal cells in two classes L5 TPC A and L5 TPC C, projected onto morphology coordinates and their respective average difference.

By contrast, the Graph–GNN model (Figure S 10, Figure 6B) reveals importance directly in the spatial domain of the neuron. Grad-CAM highlights specific nodes and branches within the morphology, thereby localizing the structural elements that drive the prediction. In correctly classified neurons, the model tends to focus on anatomically meaningful regions, such as the apical dendritic tuft. In misclassified cases, however, attention is often misdirected across competing regions or shifted toward features characteristic of another class, reflecting ambiguity in the underlying morphology.

Together, these results demonstrate that single-cell explanations provide a direct link between model predictions and interpretable morphological features. This level of analysis enables the identification of atypical or ambiguous neurons, offers a means to interrogate potential inconsistencies in expert annotations, and provides a principled framework for local quality control, such as detecting reconstruction errors or structural anomalies. More broadly, these findings highlight the value of explainability for understanding individual predictions in complex biological classification tasks.

#### 2.4.2 XAI: population-level generalization

While single-cell explanations provide detailed insight into individual predictions, an important question is whether these interpretations extend to the population level and reveal consistent class-specific features. To address this, we examined Grad-CAM patterns across groups of neurons (Figure 6C-D).

For the CNN–TMD model (Figure S 13, Figure S 14, Figure 6C), the alignment between pixel-level importance and the persistence image enables direct averaging across samples. Averaging Grad-CAM maps over correctly classified neurons yields stable patterns that closely mirror the dominant structures of the corresponding class (Figure S 13). Differences between these averaged maps highlight discriminative regions in the persistence space, providing a global description of how classes are separated in terms of their topological signatures. This consistency indicates that the CNN–TMD model captures features that are both interpretable and generalizable, and that these features can be further analyzed using established tools from topological data analysis. In the layer 5 pyramidal cell example (Figure 6C), the distinction between L5 TPC A and L5 UPC is characterized by the presence of a secondary cluster of branches in the persistence images.

In contrast, explanations derived from the Graph–GNN model exhibit greater variability at the level of individual neurons, reflecting the inherently local and sample-specific nature of morphological structure (Figure S 11, Figure S 12). Although this variability complicates direct aggregation, averaging the Grad CAM heatmats across correctly classified cells reveals recurring spatial patterns that correspond to characteristic morphological motifs (Figure S 11, Figure Figure 6D). These population-level summaries provide a complementary view to the topological analysis, capturing differences in the spatial organization of neuronal arbors. In the pyramidal cell example of layer 5 (Figure Figure 6D), the distinction between L5 TPC A and L5 UPC is characterized by the presence of an apical tuft of L5 TPC A cells that is absent from L5 UPC cells, corresponding to the topological cluster identified throughout the TMD analysis.

The interpretability of GNN-based explanations is most reliable when restricted to correctly classified samples, as misclassified neurons tend to exhibit inconsistent attention patterns. This observation underscores a fundamental distinction between the two approaches. Although CNN-TMD explanations naturally support global generalization through stable topological features, Graph–GNN explanations are particularly informative for understanding local sample-specific variability.

Together, these results demonstrate that explainability operates across scales, linking the global topological structure with the local morphological detail. The combination of CNN–TMD and Graph–GNN models therefore provides a unified and complementary framework for interpreting neuronal classification, enabling both population-level characterization and fine-grained analysis of individual cells.

## 3 Discussion

This study systematically compared multiple approaches for classifying cortical neuronal morphologies, focusing on interneurons and pyramidal cells in rodents. Three families of classifiers consistently achieved the highest performance: graph convolutional neural networks (GNNs), topological morphology descriptors (TMD), and traditional morphometrics (Figure 2). No single method dominated across all datasets, reflecting the intrinsic heterogeneity of neuronal structures. Rather than indicating limitations of individual approaches, this result underscores the multidimensional nature of neuronal morphology. Each method captures complementary aspects of neuronal organization: GNN and TMD are most effective for dendritic-based classification of pyramidal cells, while morphometrics (Ascoli et al., 2008) remain particularly informative for interneurons. Together, these findings demonstrate that robust and interpretable neuronal classification requires integrating multiple representations rather than relying on a single feature space.

Pyramidal cell classes demonstrated higher and more consistent classification accuracy than interneurons (Figure 4, Figure 5), independent of the classification method. This observation suggests that pyramidal cells exhibit more stable and distinctive morphological patterns, largely driven by the characteristic organization of their apical dendrites. In contrast, interneurons display greater morphological variability (DeFelipe et al., 2013) and are typically represented by smaller sample sizes. These factors limit classification reliability in interneurons. The improvement in performance observed when pooling interneurons across layers (Figure 2) further supports this interpretation, indicating that layer-specific distinctions are less pronounced than type-specific ones. This is consistent with transcriptomic studies showing overlapping molecular identities across layers (Gouwens et al., 2019). These results highlight the combined influence of biological variability and dataset structure on classification performance.

A central contribution of this work is the demonstration that topology and graph-based learning provide complementary and non-redundant descriptions of neuronal morphology (Figure 3). Topological data analysis, through the TMD (Kanari et al., 2018), captures *global invariants* of tree-like structures by summarizing branching hierarchies and the persistence of morphological features in a representation that is invariant to local geometric transformations. In contrast, graph neural networks (Defferrard et al., 2017) operate directly on neuronal connectivity, learning *local and hierarchical patterns* from node neighborhoods and edge relations. While topology provides a stable and scale-independent summary of structure, graph-based learning remains sensitive to fine-grained spatial and relational details. The agreement of these methods on well-defined classes, and their divergence on ambiguous ones, reflects meaningful biological variability rather than methodological inconsistency, reinforcing the value of combining both perspectives.

An essential strength of both TMD- and GNN-based approaches lies in their interpretability. By leveraging explainable artificial intelligence (XAI) techniques such as Grad-CAM, model predictions can be directly mapped to biologically meaningful features (Figure 6). In the TMD domain, pixel-level importance aligns with persistent topological structures, revealing how global branching patterns contribute to classification. In the graph domain, node-level importance localizes the contribution of specific dendritic subtrees and branching regions. Importantly, these explanations provide not only post hoc justification of predictions but also meaningful insights into the features driving model decisions. Such transparency is critical for bridging computational classification with neuroanatomical reasoning and supports the development of consensus-based labeling frameworks.

Explainability also enables analysis at multiple scales. At the level of individual neurons, XAI highlights sample-specific structural features that drive predictions, providing a powerful tool for identifying outliers, or reconstruction artifacts. This capability is particularly relevant for applications requiring fine-grained inspection, such as automated proofreading of neuronal reconstructions or quality control in large-scale datasets (Peng et al., 2015; The MICrONS Consortium et al., 2025; Shapson-Coe et al., 2024). More broadly, this level of analysis opens perspectives for personalized or case-specific investigations, where deviations from canonical morphological patterns may reflect pathological alterations or rare structural phenotypes. By localizing these deviations, XAI provides a principled framework for linking computational predictions with biologically meaningful structural variation at the level of single cells. At the population level, our findings highlight a fundamental complementarity: TMD-based explanations naturally support global generalization, whereas GNN-based explanations capture local variability and structural specificity.

Morphometrics remain valuable because they quantify physical dimensions not directly captured by structural representations, such as dendritic thickness, soma size, and tortuosity. These features are particularly informative for interneurons, where subtle geometric differences often define cell identity. However, morphometric performance depends strongly on expert-driven feature selection, which can introduce bias and limit generalizability. In contrast, TMD- and GNN-based approaches learn representations directly from morphology, reducing manual intervention and improving robustness across datasets. TMD-based models are particularly effective in low-data regimes, while GNNs offer greater flexibility and scalability across species, brain regions, and modalities.

Our results emphasize that a comprehensive understanding of neuronal morphology requires integrating multiple mathematical representations. Morphometrics, topology, and graph networks capture different aspects of neuronal structure. By combining these complementary perspectives within a unified classification and explainability framework, we obtain results that are both robust and biologically interpretable. This integrative approach also establishes a reproducible benchmark aligned with human inter-rater agreement, providing a principled upper bound for morphological classification.

The proposed framework can be extended in several directions. Incorporating additional geometric features into graph-based models, such as branch thickness or soma volume, may further improve performance. Hybrid architectures that explicitly combine topological and graph embeddings represent a promising avenue for future research. Preserving the identity of distinct neurite trees (axons, basal, and apical dendrites) could enhance both classification accuracy and interpretability. More broadly, integrating morphological, electrophysiological, and transcriptomic modalities would enable a more complete characterization of neuronal identity. In addition, we aim to explore cross-dataset generalization to leverage larger datasets for smaller sample datasets. Beyond classification, the same framework could support generative modeling of neuronal morphologies, enabling the synthesis of biologically realistic digital neurons.

In summary, topology and graph-based learning together provide a comprehensive mathematical framework for neuronal morphology. Topological descriptors capture invariant global structure, while graph-based models capture the local features that differentiate individual neurons. Their integration enables both accurate classification and interpretable insight across scales, from population-level organization to single-cell variability, offering a unified approach to understanding neuronal diversity.

## 4 Methods

### 4.1 Data collection

Experts in the field reconstructed three-dimensional neuronal morphologies, which serve as the inputs for our classification task. Patch clamped neurons were filled with biocytin and reconstructed by experts (Markram et al., 2015; Muralidhar et al., 2013). In particular, Neurolucida 360 software (Glaser and Glaser, 1990; Dickstein et al., 2016) is used to annotate points and diameters along dendritic paths and the connections between them.

Each reconstruction is a directed acyclic graph, rooted at the soma, that represents the neuron morphology embedded in the three-dimensional Euclidean space. Each vertex of this graph is associated with four floating point values – three for the coordinates (*x, y, z*) and one for the neurite diameter *d* at that point. Moreover, each vertex is assigned to either the soma or one of the neurites: the basal or apical dendrites or the axon. The connectivity between each vertex and its parent is also recorded in this representation.

### 4.2 Data Preparation

An initial sanity check commonly involves visual inspection, typically conducted by the experts who labeled the data. They identified morphologies with reconstruction artifacts, flagging those deemed insufficient in quality for morphology classification. Based on the labeled data, we performed further checks (automatic and manual) and consulted the expert labelers in cases of uncertainty.

Following the initial data quality control, the subsequent stage involved data reduction. As described in Section 2.2.1, distinct parts of neurons were found to be useful in determining morphology types, depending upon the cell type. For instance, in pyramidal cells, complete axon reconstructions were infrequent and hence unreliable as consistent features. Conversely, basal dendrites in pyramidal cells displayed minimal variation across morphology types, rendering them inadequate for morphology classification. Therefore, the distinguishing feature of pyramidal cell types is the apical dendrites. On the other hand, for interneurons, the situation is the reverse: detailed reconstructed axons provide distinctive features, while basal dendrites cannot serve as distinctive features due to their similarity across cell types. Overall, pyramidal cells were pruned to their apical dendrites, while interneurons to their axons.

Next, the selected neurites underwent simplification by removing all non-branching and non-terminating points from their morphology graphs, effectively converting their structure into binary trees, as depicted in Figure 1A. This step reduces the noise of reconstruction artifacts, due to errors arising from limitations in imaging techniques and individual scientists’ tracing styles. Another motivation for this pruning approach was to enhance compatibility with machine learning tools. Processing morphologies with only hundreds instead of thousands of vertices significantly enhances efficiency by an order of magnitude, while also removing much of the noise that obscures the important morphological features unnecessarily.

### 4.3 Feature Extraction

Traditional machine learning models such as logistic regression, support vector machines, and decision trees typically require inputs comprised of fixed-length numerical vectors. For this reason, *embedding* methods are employed to convert each input object into a fixed-length vector of numerical features. The choice of the embedding method plays a critical role since the extracted numerical features need to capture as much information as possible from the original objects to ensure the success of the machine learning classification process.

In this study, we considered two types of graph embedding methods: TMD and DeepWalk. In both approaches, each graph is transformed into a set of points in a 2-dimensional space that captures the fundamental structural properties of the raw input object. In addition, a traditional feature extraction method (Ascoli et al., 2008; Scorcioni et al., 2008) was used to extract morphological features that are typically used in classification tasks of neuronal morphologies, such as statistics on the sections lengths, number of branches, and bifurcation angles.

We also considered graph neural networks, which directly take as an input a graph object. But in this case, additional features can be relevant, as one may want to enrich the raw nodal features (e.g. the 3D coordinates of the node) with numerical features representing local or global information about the graph structure.

#### 4.3.1 Morphometrics

Traditionally neuronal morphologies have been studied based on simple comprehensive measurements that analyze different parts of its structure. A list of morphological features to study neuronal morphologies has been proposed in (Ascoli et al., 2007). Among others, features that measure the size of the neuron, such as total length and section lengths, the branching structure, such as asymmetry, and the thickness of the neuron, such as taper and the diameter power relation, were proposed as key features to describe a morphology. A long list of features has been traditionally used to describe different shapes and differentiate between them (Scorcioni et al., 2008; DeFelipe et al., 2013; Markram et al., 2015; Gouwens et al., 2019; Laturnus et al., 2020).

However, the selection of the morphological features can result in differences in the proposed classes which makes this method prone to errors and human bias. This issue arises from the fact that simple morphological features are one-dimensional projections of the neurons, a property that minimizes their complex structure and results in significant information loss. These features can be selectively tuned to assert good results and high classification accuracy based on the classification task, but are hard to generalize and difficult to interpret as a whole. We have extracted a list of morphological features that were combined with classical machine learning techniques as a comparison with previous methods (Ascoli et al., 2007; Markram et al., 2015; Gouwens et al., 2019; Laturnus et al., 2020). The extracted morphometrics include 52 features per neurite type (basal and apical dendrites and axons) and 12 global features. In total 168 features were extracted from pyramidal cells and 116 from interneurons that do not have distinct basal and apical dendrites. The complete list of morphometrics is reported in the Tables S1 and S2.

#### 4.3.2 TMD

A topological descriptor of neurons, which generates a persistence barcode from any tree-like structure was presented in (Kanari et al., 2018). The topological morphology descriptor (TMD) (Kanari et al., 2018, 2019) transforms tree-like structures, such as neuronal morphologies, that are equipped with a real-valued function *f* : *V* → ℝ on the tree nodes, to a persistence barcode through a filtration function. It produces an embedding of the graph in a two-dimensional space that has been shown to capture well the topological properties of the graph and has been used for the classification and clustering of rodent pyramidal cells (Kanari et al., 2019; Deitcher et al., 2017) and interneurons (Laturnus et al., 2020). Each branch within the tree is represented by a line in the barcode which encodes the first and the last time (in units of the function *f*) that the branch was detected in the tree structure. The persistence barcode is a collection of lines that encode the lifetime of each branch in the underlying tree structure. Equivalently, the start and end times of the branches can be represented as two-dimensional points in a persistence diagram. The two representations are equivalent and will be used interchangeably in the manuscript. An example embedding is presented in Figure S3.

#### 4.3.3 DeepWalk

DeepWalk is a popular method of graph embedding, specifically designed for learning continuous vector representations of nodes in a graph. Developed by Perozzi (Perozzi et al., 2014), DeepWalk is inspired by techniques used in natural language processing and applies them to capture the structural information of graphs. The key idea behind DeepWalk is to treat random walks on the graph as sentences and then employ skip-gram or continuous bag-of-words models to learn node embeddings. This approach enables the mapping of nodes in a graph to dense, low-dimensional vectors, preserving the inherent relationships and structural properties of the graph.

DeepWalk has applications in various domains, including social network analysis, recommendation systems, and biological network analysis. In social networks, DeepWalk can be employed to learn representations of users or nodes, capturing their relationships and interactions for tasks like link prediction or community detection. In recommendation systems, it has been used to model user-item interactions and enhance the accuracy of personalized recommendations. Additionally, in biological networks, DeepWalk has been applied to analyze protein-protein interaction networks, assisting in the identification of potential drug targets and understanding complex biological processes.

One significant advantage of DeepWalk lies in its ability to generate embeddings for nodes in large-scale graphs efficiently. By leveraging the skip-gram model and optimizing the learning process, DeepWalk can scale effectively to massive graphs, capturing both local and global structural information. The resulting embeddings can then be used for machine learning tasks, providing a compact and informative representation of the underlying graph structure, which is particularly beneficial when computational efficiency is important.

### 4.4 Classification Models

In this study, we analyzed and compared the performance of different machine learning methods for our classification task. In this section, we provide more detail about the models that we have implemented.

#### 4.4.1 Traditional Machine Learning Models

In order to establish a baseline to compare against more advanced deep learning models, we have evaluated the performance of two traditional machine learning models. More specifically, we have considered the *CART* (*classification and regression trees*) model (Lewis, 2000) using the DecisionTreeClassifier class of the scikit-learn library (Pedregosa et al., 2011b) and the *gradient-boosted trees* model (Elith et al., 2008) using the XGBClassifier class of the xgboost library (Chen and Guestrin, 2016). Both of these models are popular traditional machine learning methods based on decision trees and allow to obtain good performance with little parameter tuning. In particular, gradient-boosted trees have achieved state-of-the-art accuracy on a variety of classification tasks with structured datasets.

Simple decision trees like CART suffer from various problems like overfitting, lack of stability, or biases in imbalanced datasets. On the other hand, the gradient-boosted trees (XGBoost) algorithm addresses many of the weak points of simple decision trees and has become one of the most popular algorithms in the machine learning toolbox of many engineers and researchers.

Both of these algorithms expect a fixed-length sequence of input features. Therefore, in order to use morphological features with decision trees and XGBoost, we flatten the two-dimensional persistence images, extracted from the morphology graphs, into one-dimensional arrays of grey-scale pixel values. The morphometrics, which are also represented as fixed-length vectors, were also used as input to traditional machine learning methods.

#### 4.4.2 Convolutional Neural Networks

We argued in Section 4.3.2 that TMD is a method that captures topological properties of graph data effectively and reduces them to a two-dimensional representation in the form of barcodes, persistence diagrams (Carlsson, 2009), and persistence images (Adams et al., 2015). The persistence images, are a particularly appealing representation since it preserves the information contained in the TMD barcode and persistence diagram, but also smooths out their point-wise and discrete features into a continuous representation. This way the persistence images are insensitive to small variations in the morphology and neither are the models that take these features as inputs. This makes sense since intuitively small morphology variations should generally not change their morphological type.

Convolutional neural networks (CNN) are certainly the most natural class of machine learning models to choose from when it comes to processing image data. In the last decade, they have successfully been applied to a wide range of image-processing tasks including image classification and generation. This success is due to a range of properties that CNNs exhibit. Through the use of small two-dimensional kernels, these networks possess spatial awareness and are therefore able to recognize and learn visual patterns of the input data. At the same time, they are translationally covariant, which means that these learned patterns can be successfully recognized in any part of the picture. At this point, we refer the reader to the abundant general literature on CNNs and their applications.

Given the minor visual variance that persistence images exhibit in comparison to natural images and photographs, our goal was to choose as simple an architecture as possible. The first stage of our model is the feature extractor which consists of two convolutional layers. The second stage is the classifier which is a fully connected layer. The model inputs are 100 × 100 grey-scale persistence images. Each convolutional layer has a kernel size equal to three by three pixels, the stride of one pixel, and three output channels. Each of them is followed by a ReLU non-linearity and a max-pooling layer with both kernel and stride size equal to two. Thus after the first max-pooling the output feature map tensor has dimensions 50 × 50 × 3 and after the second one 25 × 25 × 3. Lastly, this tensor is flattened into a 1875-dimensional vector. This vector is fed into a fully connected layer with the number of output features equal to the number of classes in the respective classification task. In an alternative variant of the model, a batch-normalization layer and a ReLU non-linearity precede the fully-connected layer. Finally, a softmax activation is applied to the output of the fully connected layer to obtain a probability distribution over the classes of morphological types.

#### 4.4.3 Graph Neural Networks

Graph neural networks operate directly on the graph structure of the data. The graph data can carry different types of information: node features, edge features, node connectivity, and global features. This representation versatility can assist in faithfully capturing the underlying structures such as neurons, but at the same time presents the challenge of finding an appropriate algorithm that would make efficient use of all of these structures.

A subclass of graph neural networks is graph convolutional neural networks (Defferrard et al., 2017). This is one of the most commonly used types of graph neural networks (Graph-CNN). Similarly to the conventional convolutional neural networks, graph-CNN aggregates nearby features to derive new ones. In conventional neural networks neighboring pixels are summed over, in graph neural networks convolution happens at neighboring nodes.

The data processing pipeline within a graph convolutional neural network can be sub-divided into the same three steps as in a typical convolutional neural network: feature extraction, feature pooling, and classification.

In the feature extraction step, the graph convolution takes place and extracts node and edge features. There is a wide variety of graph convolution algorithms and layers that emerged from machine learning research. In our experiments, we used the so-called “Cheb-Conv” layer (Defferrard et al., 2017) for graph convolutions as it showed the best performance in comparison to other convolutional layers that we tried.

The goal of the feature pooling stage is to combine the extracted node, edge, and potentially global features to obtain a representation for the whole graph. Also at this point, there is a wide variety of pooling strategies that are possible.

Finally, the classification part is responsible for converting the graph embedding into a classification prediction. This can be done by standard fully connected networks and all their variations.

An extensive number of experiments were conducted to determine the best architecture and a set of configuration and training parameters. In the resulting architecture, two layers of ChebConv convolutions were used. In each of these layers, two different convolutions take place in parallel and the resulting node features are concatenated. The reason for this split is to take the graph orientation into account as best as possible. In the two parallel convolutions the graph is oriented in the opposite direction – in one the edges point from the soma towards the tips, and in the other one from the tips towards the soma. This choice has proven to produce better results than taking a single convolution on unoriented data.

Thus, overall, the model architecture of our GNN is as follows. The first two layers are graph convolutions. In the first one two ChebConv convolutions with the polynomial degree *K* = 5 are applied to the graph with one of them taking the reversed graph orientation. Each of them produces 64 feature maps. These feature maps are concatenated resulting in a total of 128 feature maps, and a ReLU non-linearity is applied. The second convolutional layer is analogous, but with each convolution producing 256 feature maps for a total of 512 feature maps, with again a ReLU non-linearity applied at the end. Next, a global pooling layer is applied. The goal of this layer is to summarise the features that are extracted for each node in the graph into a single feature vector. In our standard configuration, we used a global average pooling layer. In alternative configurations, we also tried a global sum pooling layer, as well as a learnable global attention pooling layer. One can think of the latter as a weighted sum over the graph nodes where the weights are optimized together with all other parameters of the model.

#### 4.4.4 A PersLay-Based Classifier

We showed that the neuron morphology data represented as a graph can take different forms: the original graph together with some meaningful node features, a TMD barcode, a persistence diagram, or a persistence image. In the previous sections, we described machine learning models that operate on graphs (GNNs) and on persistence images (CNNs, traditional machine learning models). In this section, we introduce a model that operates on persistence diagrams.

There are different ways in which persistence diagrams might have an advantage over graphs and persistence images. While a graph is the most faithful representation of the underlying morphology, models might struggle to capture the global properties of its structure. Indeed, GNNs that consume graphs rely on local node exploration and feature aggregation, and the graph embeddings that they produce are in the first place a summary of local neighborhoods of all of its nodes. A global view of the whole graph is never captured in a single step. Persistence diagrams and images, in contrast, do offer such a view. The points that make up the persistence diagram represent the topological structure of the whole graph. Recall that these points do not correspond to vertices in the graph, but rather represent extended topological structures. Therefore any model able to process these data would have a view on the morphology that goes beyond the local neighborhood of graph nodes. This observation holds for both persistence diagrams and images. As we explained in Section 4.3.2 the persistence image is created by applying Gaussian kernel density estimation to persistence diagrams. One may argue that this non-parametric operation is arbitrary and obscures the true nature of data. Instead one may want to embed and transform the points of a persistence diagram in a more controlled and learnable way. This is exactly what the model presented in this section sets out to do.

The main building block of the model is the PersLay embedder for persistence diagrams (Carriere et al., 2020). It takes as input persistence diagrams represented by a collection of *N* points in ℝ^2^ and produces an embedding in ℝ*^q^* for any arbitrary dimension *q* ∈ [1, ∞). More concretely our model consists of one PersLay layer with output dimension *q* = 64 that computes the morphology embeddings. It is followed by two fully connected classification layers that transform the embedding into a probability distribution over the morphology types. The first fully connected layer has an output dimension equal to 32 and is followed by a ReLU activation. The second fully connected layer’s output dimension is equal to the number of classes and is followed by a softmax activation.

How does the PersLay layer compute the embedding of a persistence diagram? The computation consists of two steps. In the first step, each point in the diagram is embedded into ℝ*^q^*by a function *f_θ_* : ℝ^2^ → ℝ*^q^*, where *θ* is the set of trainable parameters. While many choices for *f_θ_* are possible, we settled on one of the two presented in (Carriere et al., 2020), the Gaussian point transformer. In short, for *q* = 1 the embedding *f_θ_*(*x*) of a point *x* in the persistence diagram is the Gaussian distance 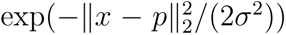, where the point *p* ∈ ℝ^2^ and width *σ* ∈ ℝ are parameters that are learnt. For a general *q* one computes *q* such distances, each with its individual parameters. For more details please see the original article. The second step is the pooling of all point embeddings into one diagram embedding. Also here many different strategies are possible as long as the pooling is invariant under the permutations of points. In our experiments, we used a simple sum of embeddings of persistence diagram points.

### 4.5 Parameter Optimization

For the results presented in Figure 2 and Figure 4C no parameter tuning was performed for any feature extraction method. Only the architectures of GNN and CNN were optimized to achieve optimal accuracy. Therefore the classification accuracy does not reflect the maximal accuracy that can be achieved by each of the reported methodologies. A sensitivity analysis of the topological approach revealed that classification accuracy can be improved from 0.8% from most layers to reach 0.9% (Figure S5b). Selecting more appropriate parameters improves significantly the classification accuracy of PersLay which was consistently performing worse with generic parameters. In addition, alternative vectorization techniques (Ali et al., 2022) were combined with TMD to test the accuracy of alternative representations instead of persistent images (Adams et al., 2015). We found that alternative vectorizations, such as entropy and life curves, achieved higher classification accuracy (Figure S6b).

### 4.6 Evaluation metrics

We performed a 10-fold cross-validation to assess the out-of-sample performance of our models, and used stratification to make sure that each split had the same distribution of labels. All metrics were computed on the out-of-sample predictions produced by the trained models on each of the 10 validation splits.

We computed several metrics to evaluate the results of the different machine learning models and feature extraction methods that we implemented. First of all, we considered *accuracy*, which is the percentage of samples for which the predicted label matches the ground truth annotation.

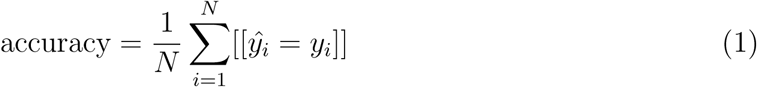

where *N* is the number of samples and [[·]] represents the indicator function, equal to 1 if the internal expression is true, and 0 otherwise.

On unbalanced datasets, it is possible to also compute *balanced accuracy*, which differs from classical accuracy in that each sample is weighted by the inverse of the percentage of samples having the same ground truth label

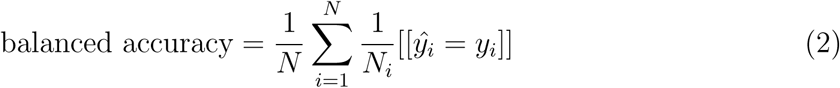

where *N_i_*is the size of the subset of samples with ground truth labels equal to *y_i_*.

As an alternative to accuracy metrics, we also considered *F1 score*, which is the harmonic mean of the precision and recall. F1 score (like precision and recall) is defined for binary classification problems, so in a multiclass classification scenario there are three strategies to compute a “global” F1 score: *micro*, *macro*, and *weighted*.

In the *micro* approach, the total number of true positives, false negatives, and false positives is considered, irrespective of the ground truth labels. In the *macro* approach, an F1 score is computed on each subset of the data having a given ground truth label, and then we take the average. The *weighted* approach is similar to the *macro*, but label imbalance is taken into account by computing the final average with weights proportional to the number of elements with that given ground truth, *N_i_*.

For our final evaluation, we needed to choose a single metric to present in our final results. However, extensive analysis of the different metrics demonstrated that there is no significant dependence in the ranking of classification methods from the selected accuracy measurement. We decided to use the accuracy score for the representation of our classification results. The main advantage of the accuracy score is that it is symmetric, i.e. accuracy(*y, ŷ*) = accuracy(*ŷ, y*). This property is useful because it allows us to compute the same metric to assess the inter-rater agreement, where we cannot consider any of the two labels provided by the expert to be more of a “ground truth” than the other, and therefore we cannot compute the other non-symmetric metrics like balanced accuracy or F1 score which require us to define which is *y* and which is *ŷ*.

#### 4.6.1 Evaluation of classification accuracy

In the majority of machine learning literature, classifiers are typically designed with the primary goal of surpassing existing models. However, the intricacies of neural grouping in the context of biological systems have long been a subject of debate. In a seminal work by DeFelipe et al. (DeFelipe et al., 2013), it was elucidated that expert classifications exhibit a notable bias contingent upon the reconstructor executing the task. Consequently, the pursuit of attaining 100% accuracy in delineating specific neuronal groupings is not a realistic expectation, and may potentially signify overfitting to the inherent noise within biological datasets.

To circumvent an overreliance on the noise prevalent in biological data for achieving higher classification scores, we advocate for the inclusion of two pivotal components in the assessment criteria for classifiers. Firstly, emphasis should be placed on achieving optimal classification accuracy (or minimizing error) while accounting for inter-rater agreement. Secondly, we postulate that the elucidation of classification results is equally crucial, thereby demonstrating the significance of explainability in the context of neuronal grouping.

To evaluate the performance of different classification methods, we performed two baseline comparative scores: the chance level and the expert agreement.

Most of the datasets were generated within the LNMC lab, and therefore an independent classification could not be performed. For the external dataset (Janelia) manual classification was performed by two experts and their results were compared to assess inter-rater agreement.

#### 4.6.2 Random score computation

We computed a baseline accuracy score, to compare it against the performance of our models. In particular, we considered the average performance of “chance agreement” for a random model that predicts the value of each label independently from a categorical distribution having weights equal to the observed label frequencies.

The probability of any given sample to have label *k* (for *k* in 1*…K*) is

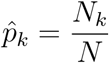

If we consider assigning random labels according to the observed occurrences, this means that the predicted label of the i-th sample, *ŷ_i_* is given by

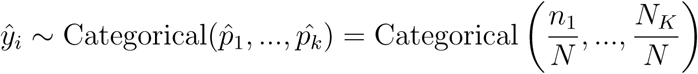

Then, the probability of the event *y_i_* = *ŷ_i_* is computed using the Law of Total Probability as

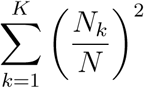

### 4.7 Inter-rater agreement

As mentioned above, (unbalanced) accuracy was used as our evaluation metric for several reasons, notably its symmetry, allowing for the computation of an inter-rater agreement score. This score can then be directly compared to the performance of our machine learning models, providing valuable insights into their efficacy. Specifically, we collected annotations from two distinct experts regarding the Janelia pyramidal cells of layer 5. Based on these annotations, we computed the inter-rater agreement by the number of labels that are shared between the two experts compared to the total number of expert evaluations.

### 4.8 Explainable Artificial Intelligence (XAI)

The use of Grad-CAM (Gradient-weighted Class Activation Mapping) as an explainable AI technique provides valuable insights into the decision-making process of both CNN and GNN models.

For CNNs, which operate on image data, Grad-CAM highlights the areas of the image that contribute more to the decision of the label for each sample. This allows researchers to visually inspect which regions of the input image are most influential in the classification process. By analyzing these highlighted areas, one can gain an understanding of the specific features or patterns that the CNN model is focusing on to make its predictions. This method of explanation is particularly useful for tasks such as image classification, where the input data consists of visual information.

On the other hand, for GNNs, which operate on graph-structured data, Grad-CAM outputs the importance of each node in the graph structure to the decision of the labels. This means that instead of highlighting regions in an image, Grad-CAM for GNNs identifies the specific nodes within the graph representation of the data that are most influential in determining the classification outcome. By examining the importance scores assigned to each node, researchers can understand which parts of the graph structure are most informative for the GNN model’s decision-making process. This approach provides insights into how the connectivity and topology of the graph contribute to the classification results, offering a deeper understanding of the underlying relationships within the data.

In summary, while Grad-CAM serves as an explainable AI technique for both CNN and GNN models, its interpretation varies based on the nature of the input data. For CNNs, Grad-CAM highlights important image regions, whereas for GNNs, it identifies influential nodes in the graph structure. This distinction allows researchers to gain meaningful insights into the inner workings of each model and the factors driving their classification decisions.

In addition to advanced XAI techniques, we compute feature importance with XG-Boost (Adler and Painsky, 2021) for the morphometrics to assess the contribution of different morphological features in the classification results (Figure S2). More specifically, we rank the morphological features according to their weight, which counts the number of times a feature is used to split the data across all XGBoost trees. In Figure S2 we present the 10 most significant features for each classification task (A: interneurons, B: pyramidal cells).

## 5 Acknowledgements

The authors thank Prof. Kathryn Hess for her useful contributions to the discussions on topological descriptors, Prof Pierre Vandergheynst for the useful feedback and the suggestion of the deepwalk model for comparison, and Jan Krepl for his useful contributions to the machine learning techniques. We thank LNMC for the reconstructions and the experimental work (Data collectors: Rodrigo de Campos Perin, Shruti Muralidhar, Thomas Berger) that made this study possible.

## 7 Author contributions

L.K. and F.C. conceived the study and supervised the experiments. S.S., E.D., L.K., F.C. developed the code and conducted the computational experiments. M.D. designed machine learning architectures for graph neural networks and supervised the study. J.B. and T.N. contributed to the code development and performed computational experiments. Y.S. conducted experiments, reconstructed neurons, and carried out expert classification. J.M. conducted expert classification. F.S. and H.M. secured funding and supervised the study. All authors contributed to writing and editing the manuscript throughout the process.

## 8 Data availability

The data used for this paper are available from (Kanari et al., 2022) in https://zenodo.org/records/5909613 and from (Winnubst et al., 2019) in http://ml-neuronbrowser.janelia.org/.

## 9 Code availability

The code used for this paper is openly available in our “morphoclass” repository on GitHub at https://github.com/BlueBrain/morphoclass. The dataset and scripts used for this paper will be made available upon publication. An example is given in Example of code use.

