## Supplementary Information for "An interpretable deep learning framework for classifying neuronal morphologies using topology and graph neural networks"

### Supporting information

#### Interneuron cell types

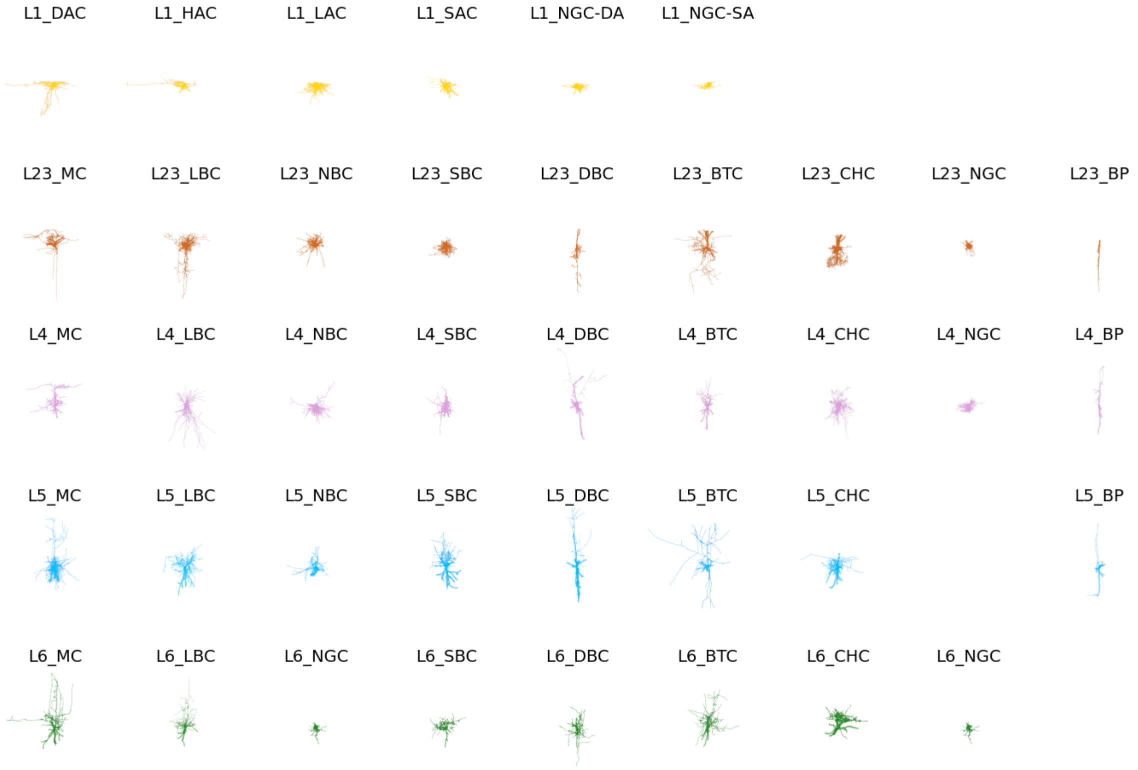


Figure S 1: **Interneuron cell types.** Examples of all interneuron cell types from cortical layers 1 (yellow), 2 and 3 (orange), 4 (purple), 5 (blue) and 6 (green). For details about the cell types see Table S4.

#### List of Morphometrics

| | |
|---|---|
| mean local bifurcation angles | mean section radial distances |
| mean number of bifurcations | max section Strahler orders |
| mean number of leaves | mean section Strahler orders |
| mean number of neurites | min section term lengths |
| mean number of sections | max section term lengths |
| mean number of segments | mean section term lengths |
| mean partition asymmetry | mean section term radial distances |
| mean partition asymmetry length | mean section tortuosity |
| max principal direction extents | mean section volumes |
| mean remote bifurcation angles | mean segment meander angles |
| mean section areas | mean terminal path lengths |
| min section bif lengths | mean total area |
| max section bif lengths | sum total area |
| mean section bif lengths | min total area per neurite |
| mean section bif radial distances | max total area per neurite |
| min section branch orders | mean total area per neurite |
| max section branch orders | mean total length |
| mean section branch orders | sum total length |
| mean section end distances | min total length per neurite |
| min section lengths | max total length per neurite |
| max section lengths | mean total length per neurite |
| mean section lengths | mean total volume |
| min section path distances | sum total volume |
| max section path distances | min total volume per neurite |
| mean section path distances | max total volume per neurite |
| median section path distances | mean total volume per neurite |

Table S 1: List of morphometric features per neurite type (basal and apical dendrite and axon).

| |
|---|
| mean max radial distance |
| mean number of neurites |
| mean number of sections per neurite |
| mean soma radius |
| mean soma surface area |
| mean soma volume |
| mean total area per neurite |
| mean total depth |
| mean total height |
| mean total length per neurite |
| mean total volume per neurite |
| mean total width |

Table S 2: List of morphometrics per neuron.

| Acronym | Cell type name |
| --- | --- |
| TPC | Tufted pyramidal cell |
| TPC:A | Large tufted pyramidal cell |
| TPC:B | Early bifurcating tufted pyramidal cell |
| TPC:C | Small tufted pyramidal cell |
| UPC | Untufted pyramidal cell |
| SSC | Spiny stellate cell |
| IPC | Inverted pyramidal cell |
| BPC | Bitufted pyramidal cell |
| HPC | Horizontal pyramidal cell |

Table S 3: List of pyramidal cell types

| Acronym | Cell type name |
| --- | --- |
| MC | Martinotti cell |
| LBC | Large basket cell |
| NBC | Nest basket cell |
| SBC | Small basket cell |
| DBC | Double bouquet cell |
| BTC | Bitufted cell |
| CHC | Chandelier cell |
| NGC | Neurogliaform cell |
| BP | Bipolar cell |
| DAC | Descending axon cell |
| HAC | Horizontal axon cell |
| LAC | Large axon cell |
| NGC-DA | Neurogliaform cell with dense axonal arbor |
| NGC-SA | Neurogliaform cell with sparse axonal arbors |
| SAC | Small axon cell |

Table S 4: List of interneuron cell types

#### Feature importance

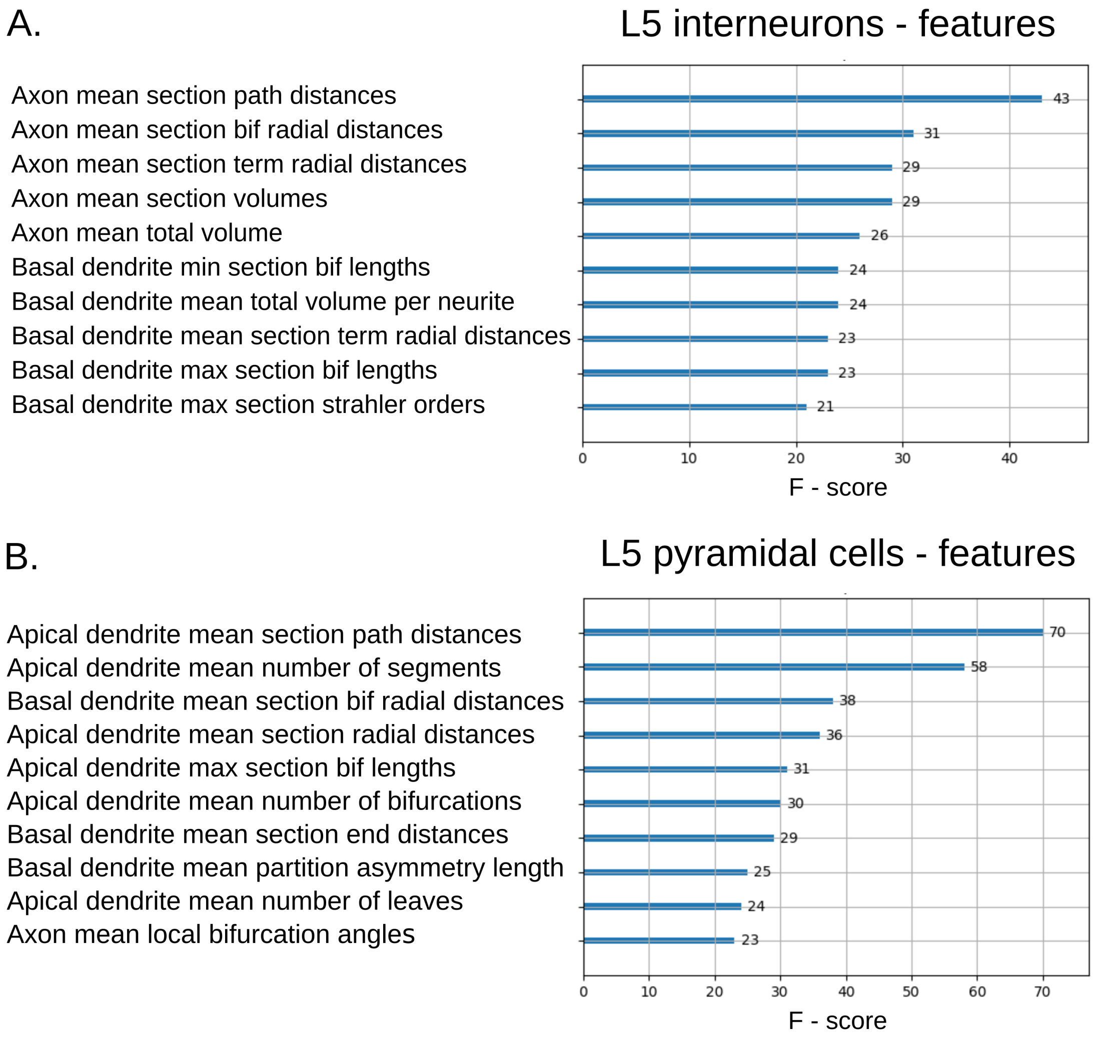


Figure S 2: **Morphometrics feature importance for layer 5 interneurons and pyramidal cells.** Feature importance is computed using XGBoost weights, defined as the number of times each feature is used to split the data across all trees in the model. (A) For interneurons, the most important features are predominantly axonal. (B) For pyramidal cells, the most important features are mainly associated with apical dendrites.

#### Topological Morphology Descriptor

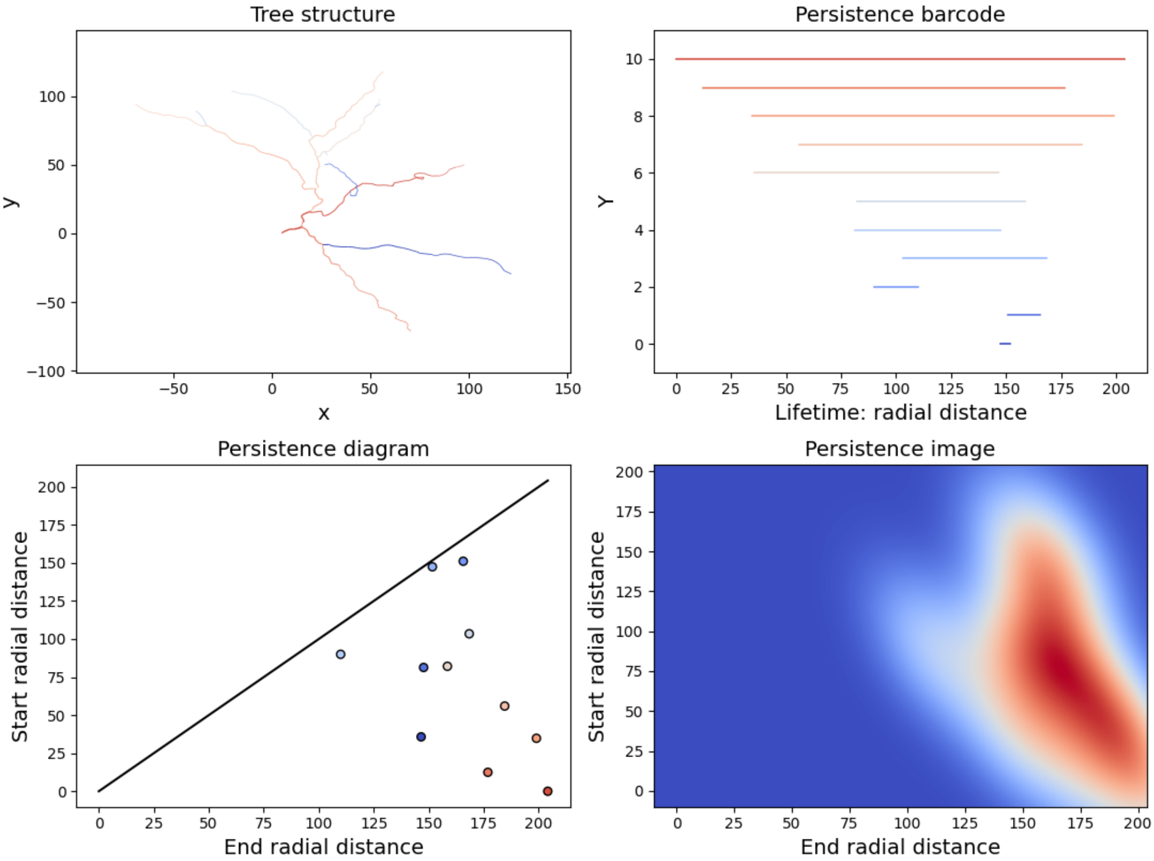


Figure S 3: **Topological Morphology Descriptor extracted from a dendritic tree.** (A). The reconstructed tree. Colors represent the branches and the bars in the persistence barcode, ranging from longer (red) to smaller branches (blue). (B). The TMD barcode produced through filtration of the apical graph using radial distances. (C). The persistence diagram presents the same information plotted in two dimensional space where axes represent the birth and the death of each segment of the bar code. (D). The persistence image shows a Gaussian kernel density estimate applied to the persistence diagram in C.

#### Detailed classification results

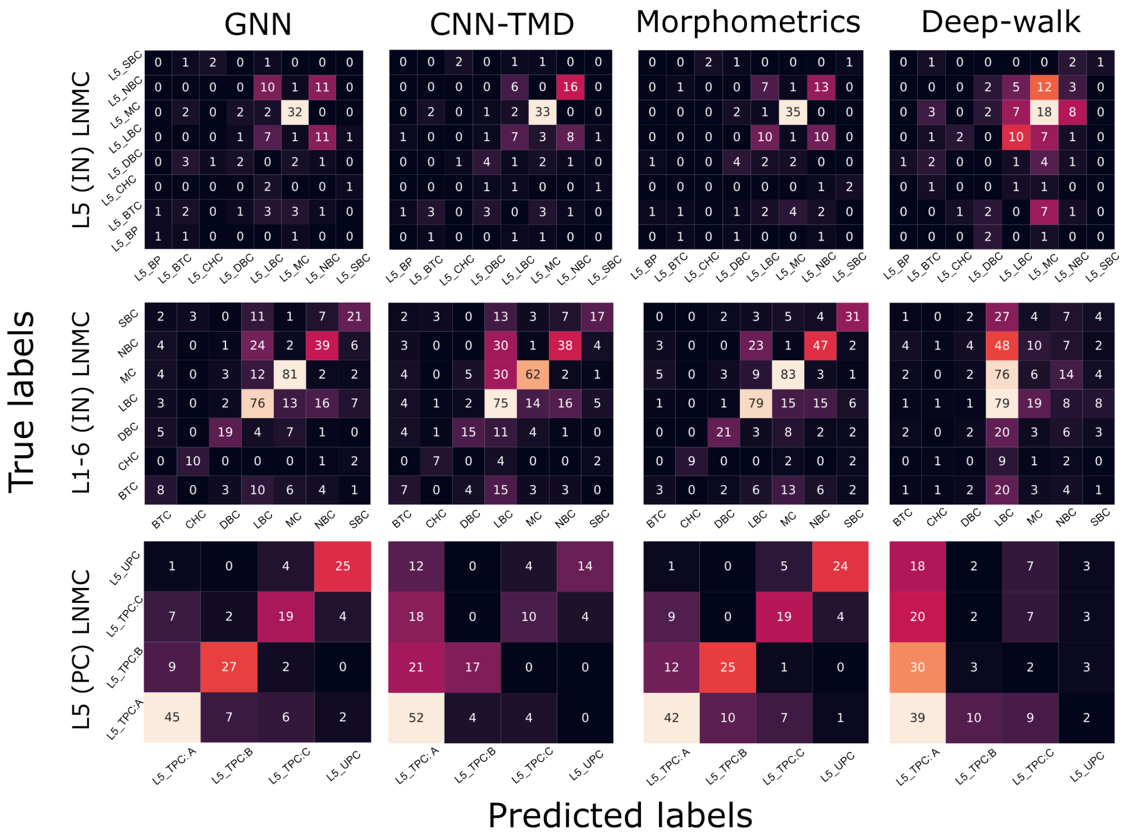


Figure S 4: **Confusion matrices examples.** Confusion matrices for different classification methods (columns) and datasets (rows). The accuracy of each classification method is evaluated by comparing the true labels to the predicted labels. The color map illustrates the number of predictions (black: lowest number of samples, white: highest number of samples per experiment). Perfect accuracy (all predicted labels correspond to true labels) is indicated by white on the diagonal. The colorscale (dark:minimum number to white:maximum number) is not normalized across different data.

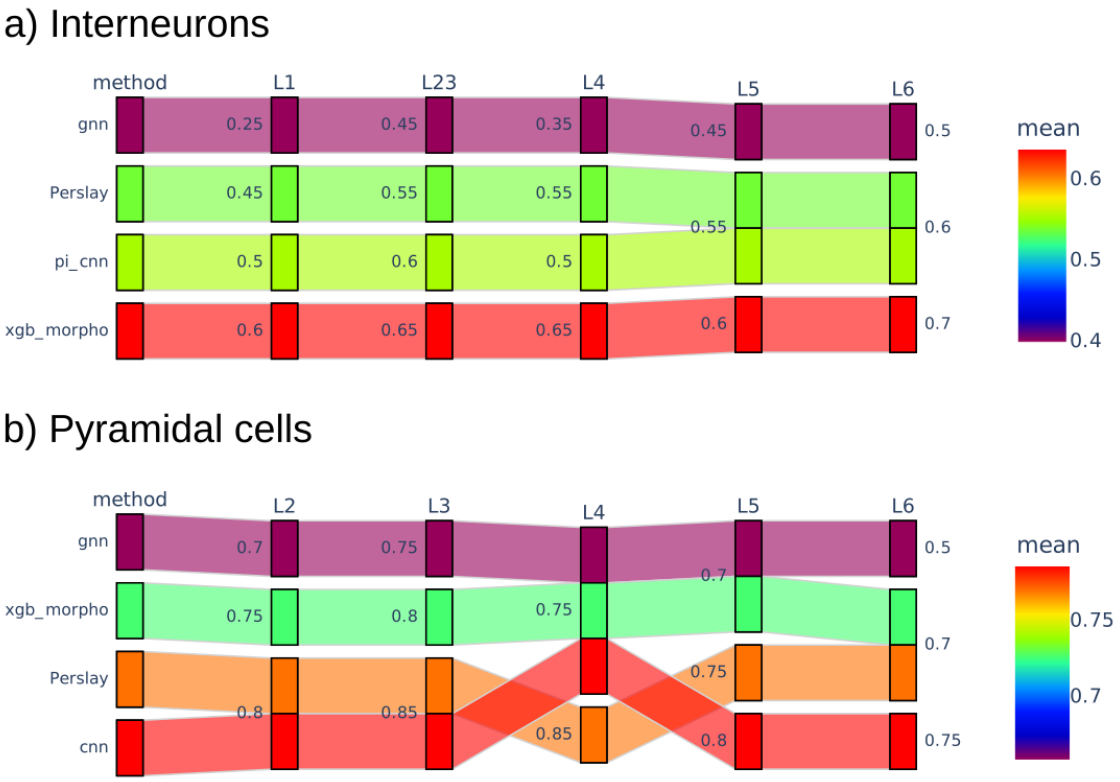


Figure S 5: **Classification accuracy with fine-tuned parameters across cortical layers.** Sankey-style plots show the classification accuracy of different methods, including PersLay with optimized parameters, across cortical layers for (a) interneurons and (b) pyramidal cells. Each row corresponds to a classification method, and columns represent cortical layers. Colors encode the mean accuracy, as indicated by the color bar, with warmer colors representing higher performance. Numerical values indicate the accuracy achieved by each method within each layer.

| Scores | All | AIBS | Petilla | Laturnus |
|---|---|---|---|---|
| Accuracy | 0.69 | 0.69 | 0.47 | 0.58 |
| Macro | 0.69 | 0.69 | 0.46 | 0.60 |
| Weighted | 0.69 | 0.69 | 0.47 | 0.58 |

Table S 5: **Classification accuracy using alternative morphometric feature sets for layer 5 pyramidal cells.** Classification performance is reported for different subsets of morphometric features, compared to the full feature set. Results indicate that alternative feature selections generally achieve comparable or lower accuracy than the complete set of morphometrics, suggesting that no reduced subset consistently outperforms the full representation. This supports the use of comprehensive morphometric descriptors to capture the full variability of neuronal morphology.

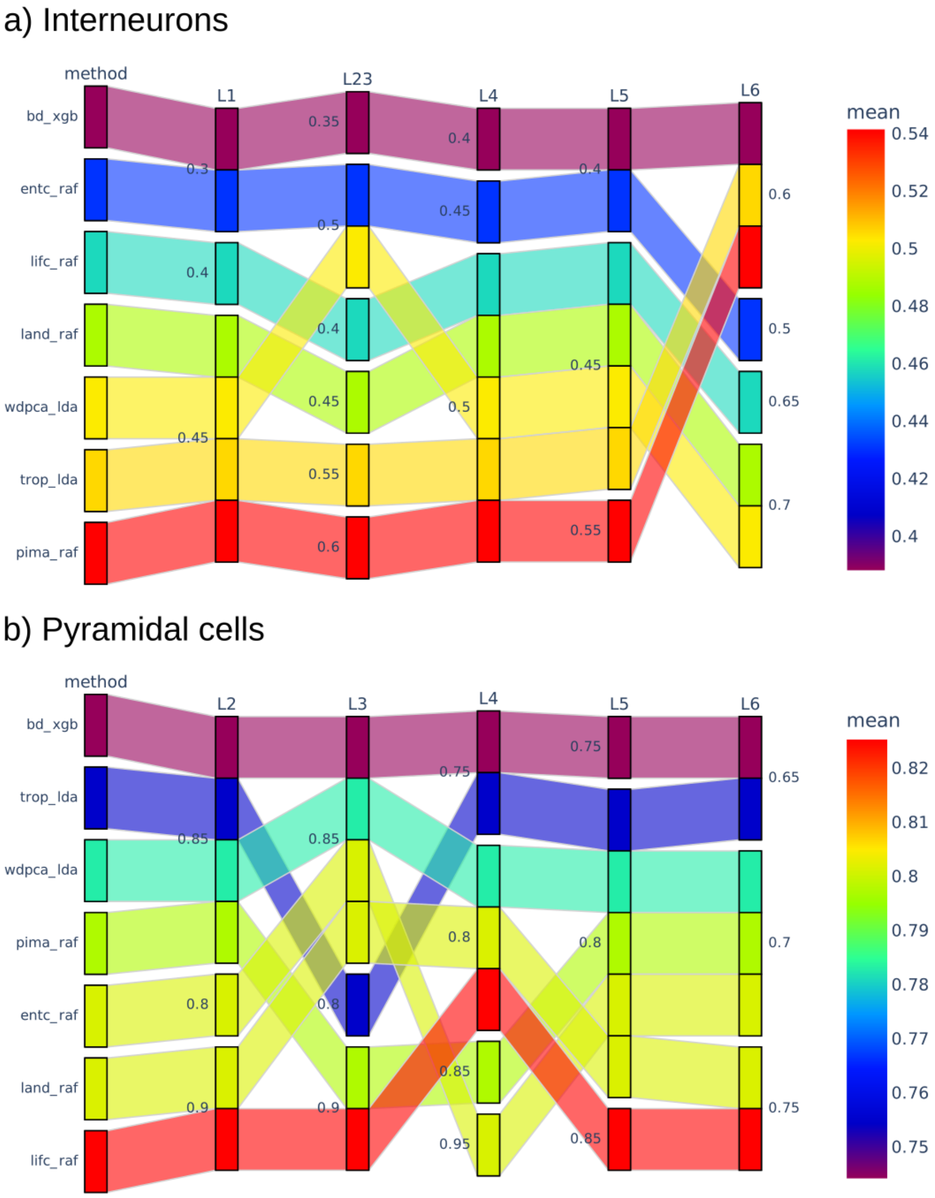


Figure S 6: **Classification accuracy across topological vectorization methods and cortical layers.** Sankey-style plots show the classification accuracy of different topological vectorization methods across cortical layers for (a) interneurons and (b) pyramidal cells. Each row corresponds to a method, and columns represent cortical layers. Colors encode the mean classification accuracy, as indicated by the color bar, with warmer colors representing higher performance. Numerical values denote the accuracy achieved by each method within each layer.

| Dataset | Method | Basal dendrites | Axons | Apical dendrites | All |
|---|---|---|---|---|---|
| L5-in | GNN | 0.38 | 0.50 | - | 0.54 |
| L5-in | TMD-CNN | 0.33 | 0.57 | - | 0.55 |
| L5-in | TMD-XGB | 0.32 | 0.50 | - | 0.49 |
| L5-pc | GNN | 0.36 | 0.36 | 0.73 | 0.48 |
| L5-pc | TMD-CNN | 0.39 | 0.33 | 0.58 | 0.45 |
| L5-pc | TMD-XGB | 0.30 | 0.28 | 0.75 | 0.59 |

Table S 6: **Classification accuracy across alternative neurite types.** Classification performance is reported for different neurite-specific representations, including basal dendrites, apical dendrites, axons, and their combination, for both layer 5 interneurons and pyramidal cells. The results highlight that axonal features provide the most discriminative information for interneuron classification, whereas apical dendrites are most informative for pyramidal cell classification. In some cases, combining all neurite types further improves performance, particularly for interneurons, suggesting that complementary structural information across neurite compartments can enhance classification accuracy.

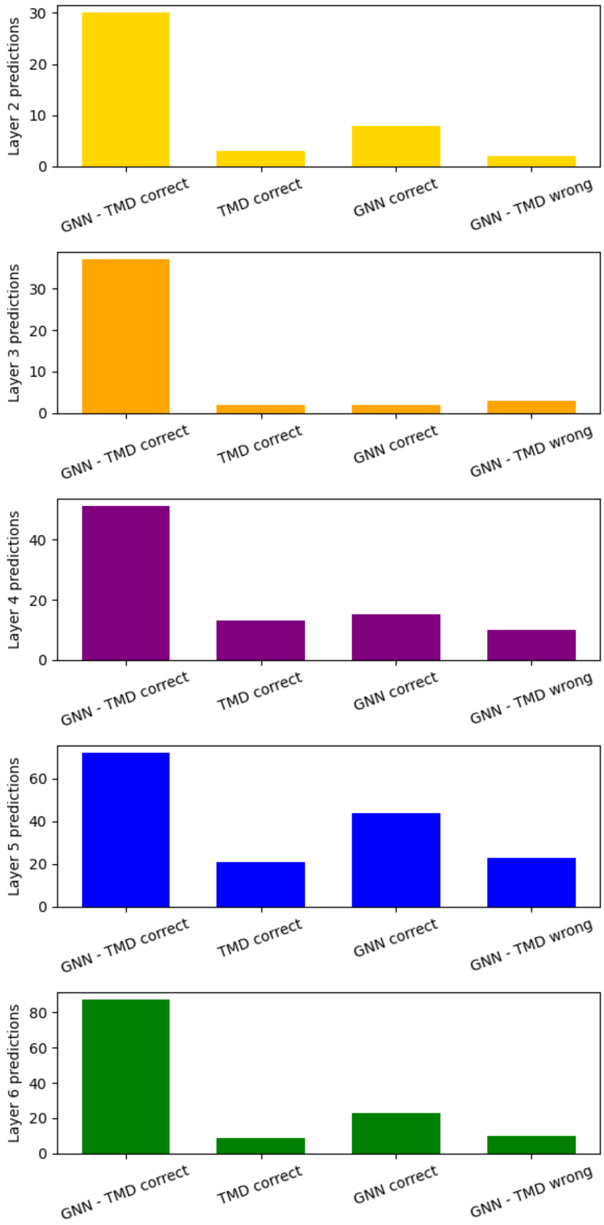


Figure S 7: **Complementarity between TMD-CNN and Graph-GNN models across cortical layers.** Bar plots show the distribution of prediction outcomes for pyramidal cells in layers 2–6, comparing graph-based (GNN) and topology-based (TMD) classification methods. For each layer, predictions are partitioned into four categories: correctly classified by both methods ("GNN–TMD correct"), correctly classified only by TMD-CNN ("TMD correct"), correctly classified only by GNN ("GNN correct"), and misclassified by both methods ("GNN–TMD wrong").

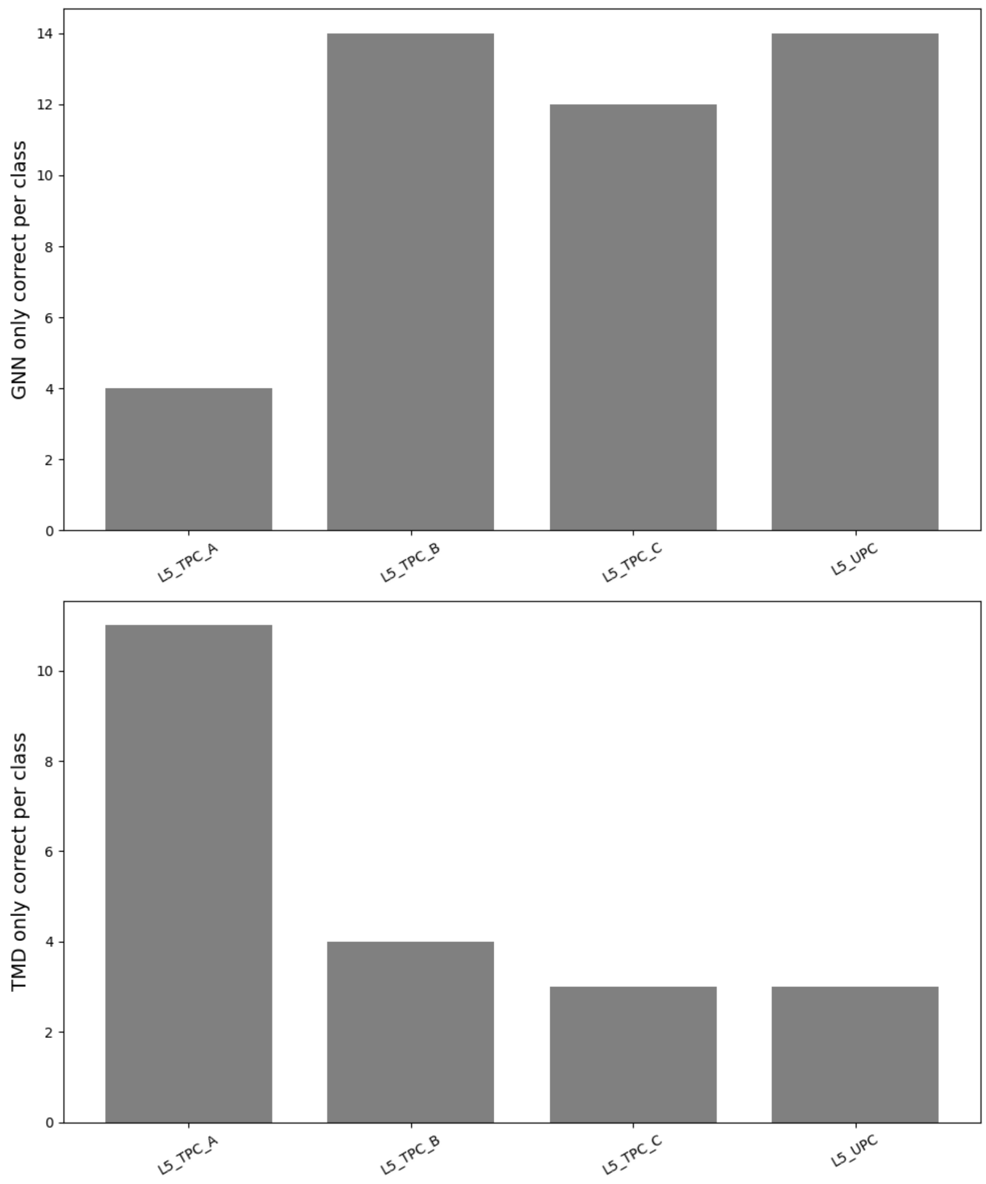


Figure S 8: **Class-specific complementarity between TMD-CNN and Graph-GNN models in layer 5 pyramidal cells.** Bar plots show the number of correctly classified samples per class that are uniquely identified by each method. The top panel ("GNN only correct per class") represents neurons correctly classified by the graph-based GNN model but misclassified by the TMD-CNN. The bottom panel ("TMD only correct per class") shows neurons correctly classified by the TMD-CNN but misclassified by the GNN.

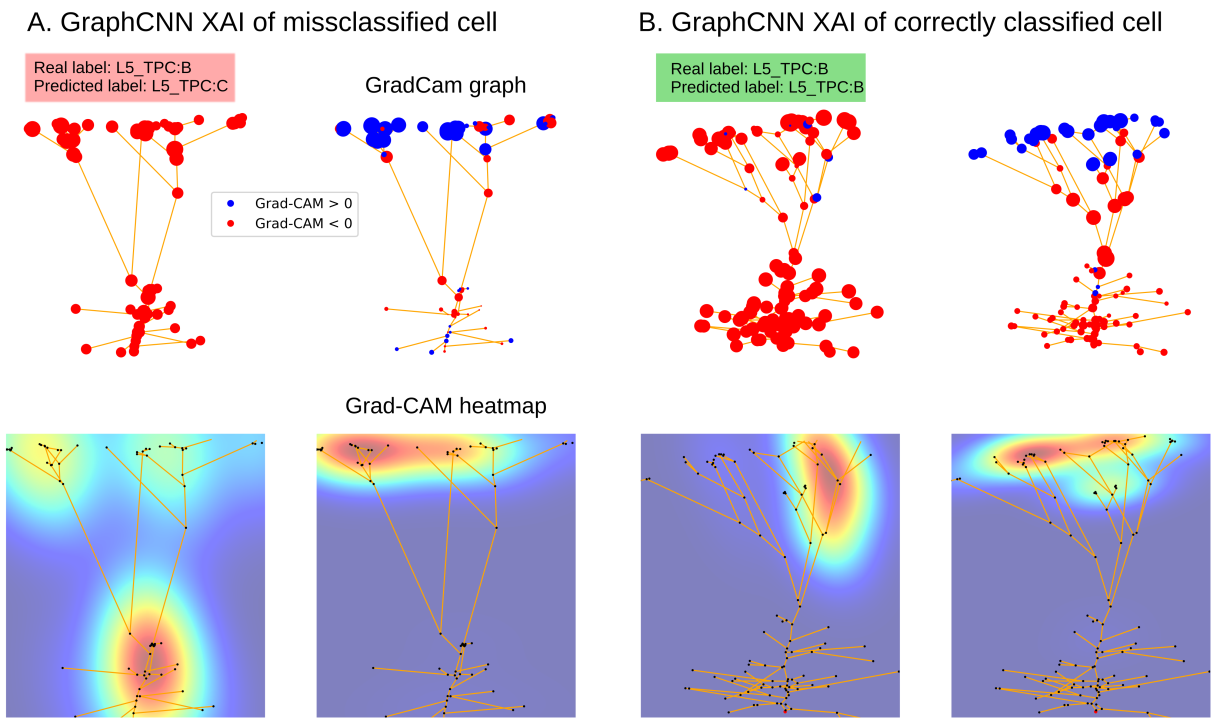


Figure S 9: **Grad-CAM-based explainability for Graph-GNN predictions.** (A) Example of a misclassified neuron (true label: L5 TPC:B). (B) Example of a correctly classified neuron (true label: L5 TPC:B). The top row shows node-level importance on the graph, where node size reflects the magnitude of the Grad-CAM signal and color indicates its sign (blue: positive contribution to the predicted class; red: negative contribution). The bottom row presents spatial heatmaps of the aggregated Grad-CAM signal, illustrating the distribution of model attention across the neuronal morphology, from high attention (red) to low attention (blue).

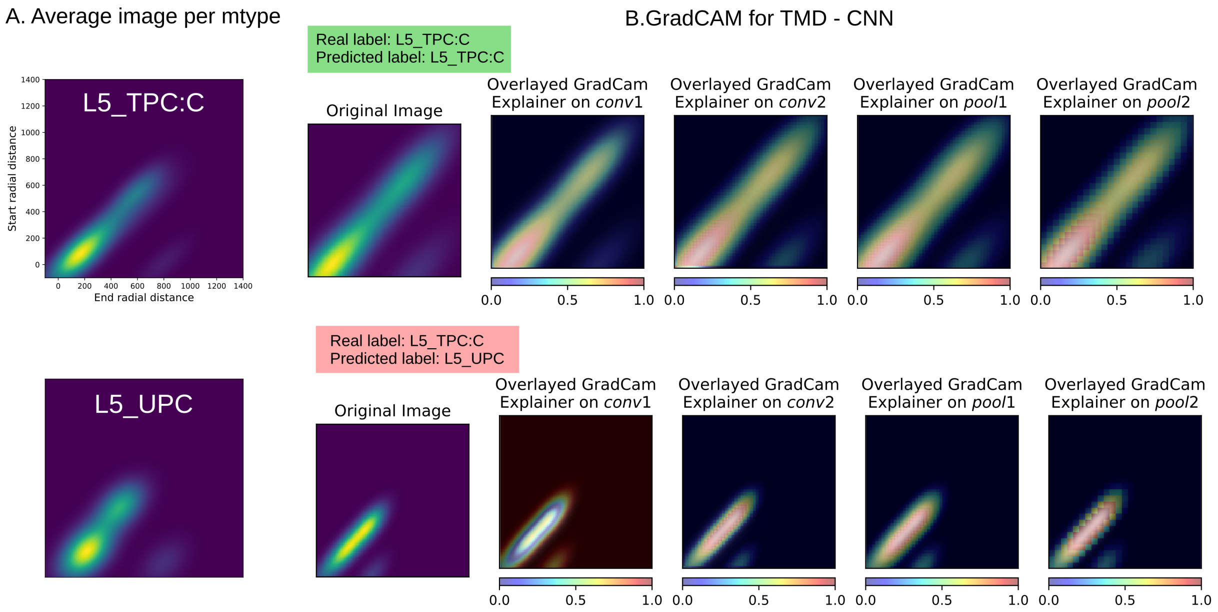


Figure S 10: **Grad-CAM-based explainability for TMD-CNN predictions.** (A) Example of a correctly classified neuron (true label: L5 TPC:C). (B) Example of a misclassified neuron (true label: L5 TPC:C, predicted: L5 UPC). The left panels show the average topological images per m-type and the corresponding input image. The right panels display Grad-CAM visualizations at different network layers (conv1, conv2, pool1, pool2), highlighting pixel-level contributions to the model's prediction. Color intensity represents the magnitude of attention, ranging from low (blue) to high (yellow/red).

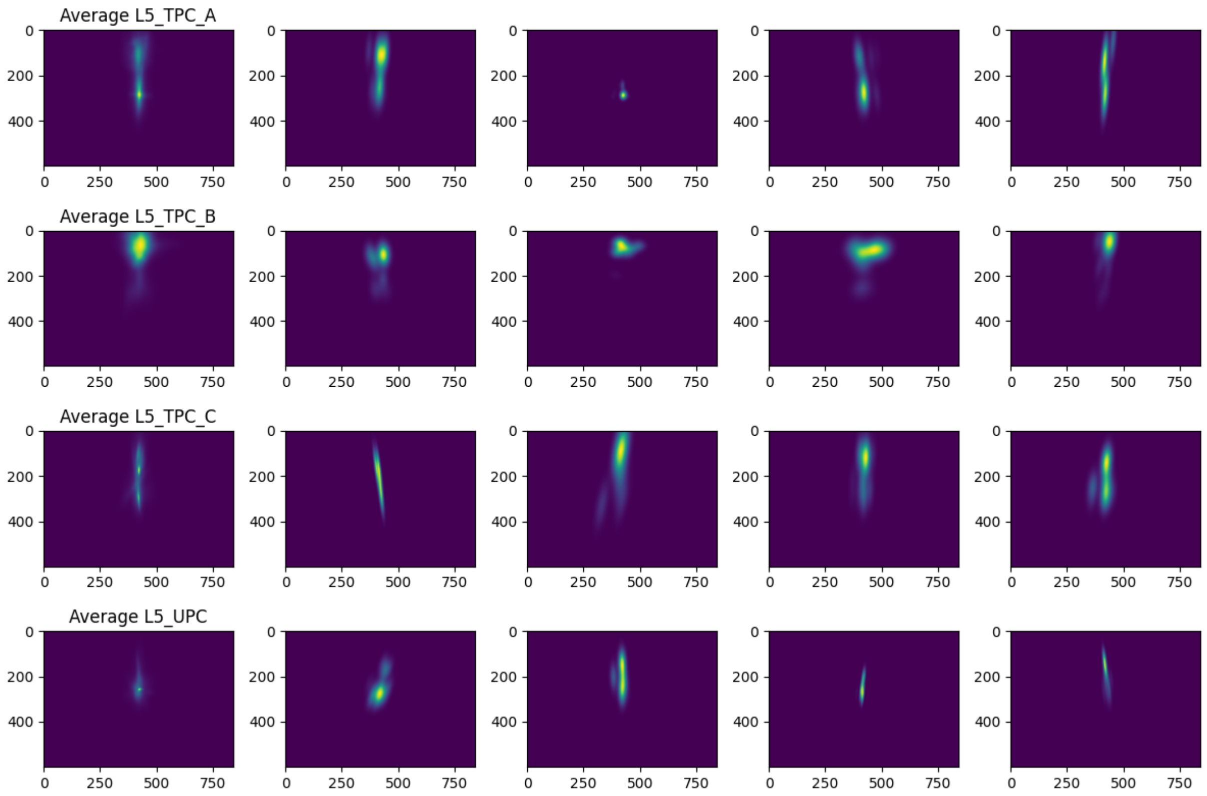


Figure S 11: **Population summary of GNN explainability for correctly classified layer 5 pyramidal cell types.** The figure shows the average Grad-CAM heatmaps obtained from the correct predictions of the GNN model for four layer 5 pyramidal cell classes: L5_TPC:A, L5_TPC:B, L5_TPC:C, and L5_UPC. Each row corresponds to one class. The first panel within a row shows a representative averaged saliency map, the rest of the panels show a random example within that class. Brighter regions indicate higher model attention, whereas darker regions indicate lower contribution to the classification decision.

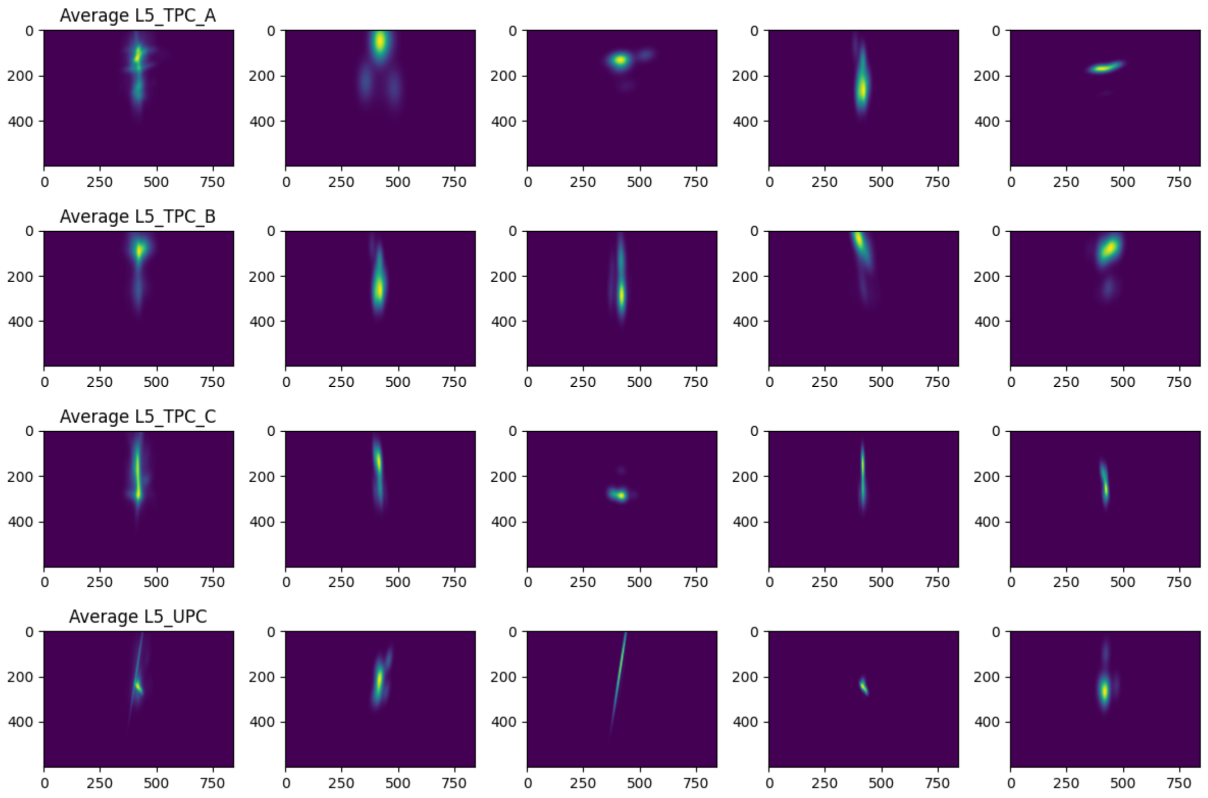


Figure S 12: **Population summary of GNN explainability for wrongly classified layer 5 pyramidal cell types.** The figure shows the average Grad-CAM heatmaps obtained from the wrong predictions of the GNN model for four layer 5 pyramidal cell classes: L5_TPC:A, L5_TPC:B, L5_TPC:C, and L5_UPC. Each row corresponds to one class. The first panel within a row shows a representative averaged saliency map, the rest of the panels show a random example within that class. Brighter regions indicate higher model attention, whereas darker regions indicate lower contribution to the classification decision.

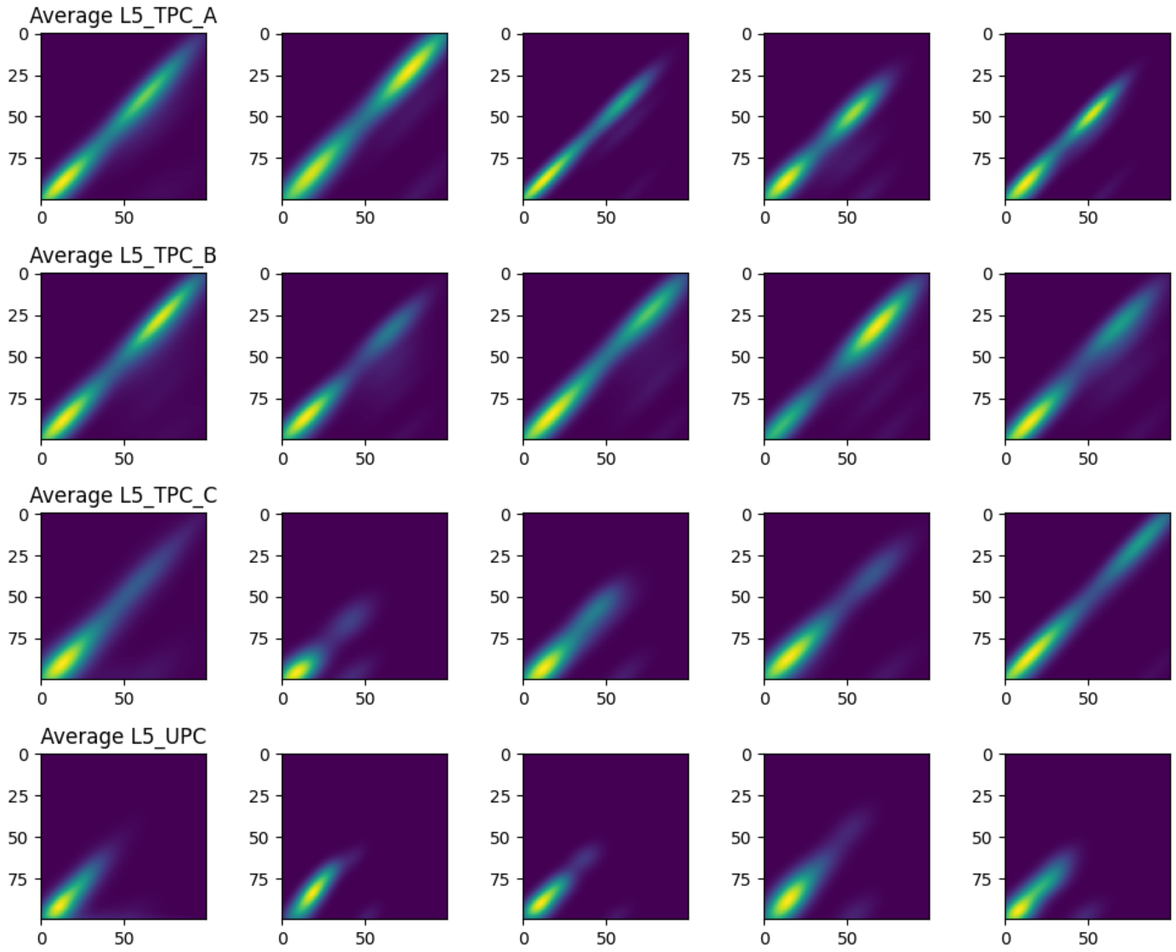


Figure S 13: **Population summary of CNN TMD explainability for correctly classified layer 5 pyramidal cell types.** The figure shows the average Grad-CAM heatmaps obtained from the correct predictions of the CNN TMD model for four layer 5 pyramidal cell classes: L5_TPC:A, L5_TPC:B, L5_TPC:C, and L5_UPC. Each row corresponds to one class. The first panel within a row shows a representative averaged saliency map, the rest of the panels show a random example within that class. Brighter regions indicate higher model attention, whereas darker regions indicate lower contribution to the classification decision.

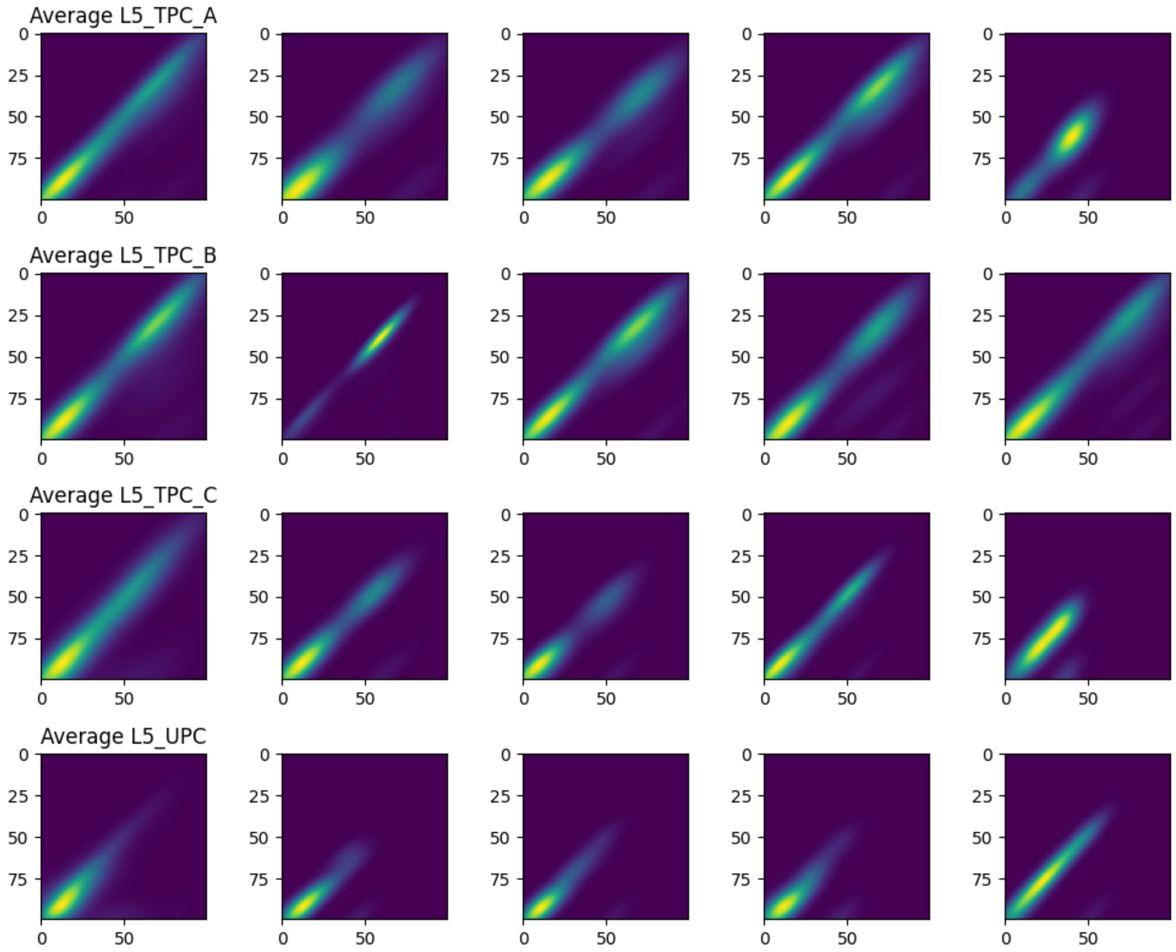


Figure S 14: **Population summary of CNN - TMD explainability for wrongly classified layer 5 pyramidal cell types.** The figure shows the average Grad-CAM heatmaps obtained from the wrong predictions of the CNN - TMD model for four layer 5 pyramidal cell classes: L5_TPC:A, L5_TPC:B, L5_TPC:C, and L5_UPC. Each row corresponds to one class. The first panel within a row shows a representative averaged saliency map, the rest of the panels show a random example within that class. Brighter regions indicate higher model attention, whereas darker regions indicate lower contribution to the classification decision.

##### Example of code use

We present below a basic example to demonstrate the simple use of morphoclass package. More details are provided in https://github.com/BlueBrain/morphoclass/.

Preprocessing:

```
morphoclass preprocess-dataset
       --dataset-type pc-L5
       --morphologies-dir data/raw/pc-L5/morphologies
       --db-file data/raw/pc-L5/neurondb.dat
       --output-csv-path data/preprocessed-pc-L5.csv
       --output-report-path reports/preprocess-dataset/pc-L5.html
```

Feature extraction:

```
$ morphoclass extract-features
     data/pyramidal-cells/L5/dataset.csv
     apical
     image-tmd-rd
     extract-features/pc-L5/apical/image-tmd-rd/
```

Training:

```
$ morphoclass train
     --features-dir  extract-features/pc-L5/apical/image-tmd-rd/
     --model-config  training/configs/model-xgb.yaml
     --splitter-config training/configs/splitter-stratified-k-fold.yaml
     --checkpoint-dir training/pc-L5-apical-image-tmd-rd-xgb/
```

Evaluation:

```
$ morphoclass evaluate performance
     training/pc-L5-apical-image-tmd-rd-xgb/checkpoint.chk
     evaluation_report.html
```
